# An isogenic human stem cell model reveals NGLY1-dependent control of microglial development

**DOI:** 10.64898/2026.11.13.766515

**Authors:** Haiwei Zhang, Haipeng Xue, Mengmeng Jin, Ting Wang, Adam J. Rauckhorst, Peng Jiang, Yu-Chieh Wang, Ying Liu

## Abstract

NGLY1 is a cytosolic N-glycanase involved in glycoprotein quality control and proteasome-mediated protein degradation and is critical for normal brain development and function. Although neuronal alterations caused by NGLY1 deficiency have been extensively characterized in animal and human cell models, how NGLY1 deficiency affects human glial cells, particularly microglia, the brain-resident innate immune cells, remains largely unknown. Using patient-derived NGLY1-deficient human induced pluripotent stem cell (hiPSC) lines and their corresponding CRISPR-mediated gene-corrected isogenic lines, we found that NGLY1-deficient primitive macrophage progenitors (PMPs) exhibited increased GlcNAc-Asn (GNA) accumulation and elevated expression of RUNX1, a key transcription factor regulating myeloid lineage development, accompanied by abnormal proliferation. Following further differentiation, NGLY1-deficient human microglia exhibited reduced morphological complexity and impaired phagocytic capacity. In addition, NGLY1-deficient microglia showed increased expression of pro-inflammatory cytokines, including IL-1β, IL-6, and TNFα, under both basal and stimulated conditions, indicative of a heightened inflammatory state. Mechanistically, our data showed that RUNX1 upregulation induced by NGLY1 deficiency partially mediated impaired PMP proliferation and subsequent microglial abnormalities, implicating RUNX1 dysregulation in NGLY1-associated microglial dysfunction and identifying RUNX1 as a potential therapeutic target. Together, using patient-derived and gene-corrected isogenic hiPSCs, our study establishes a human stem cell platform for modeling NGLY1 deficiency and reveals that impaired deglycosylation alters human microglial developmental trajectories and ultimately compromises microglial homeostasis and function. These microglial abnormalities may contribute to the neurological deficits observed in patients carrying loss-of-function variants in NGLY1.

## Introduction

*NGLY1* encodes a cytosolic N-glycanase that removes N-linked glycans from misfolded glycoproteins during protein quality-control pathways (Caglayan et al., 2015; Enns et al., 2014; Huang et al., 2015; Need et al., 2012; Suzuki, 2015). Loss-of-function variants in *NGLY1* cause an ultra-rare congenital disorder associated with developmental delay, movement abnormalities, seizures, liver abnormalities, alacrima or hypolacrima, and peripheral neuropathy (Enns *et al*., 2014; Lam et al., 2017; Need *et al*., 2012; Tong et al., 2023; Zhang et al., 2025). NGLY1 has also emerged as a key regulator of proteostasis-linked transcriptional programs. For example, inhibition or loss of NGLY1 function impairs the post-translational processing and activation of NRF1/NFE2L1, thereby compromising NRF1/NFE2L1-mediated proteasome homeostasis and ultimately disrupting cellular proteostasis (Lehrbach et al., 2019; Mueller et al., 2020; Tomlin et al., 2017; Yang et al., 2018). Recent animal studies further revealed the complexity of NGLY1-associated neuropathology, showing that NGLY1 deficiency affects multiple aspects of nervous system development and function, including neuronal differentiation and maturation, axonal integrity, synaptic function, and neural circuit organization, and can lead to progressive neurological and behavioral abnormalities (Asahina et al., 2021a; Asahina et al., 2021b; Asahina et al., 2020; Du et al., 2024; Mesika et al., 2022; Yang et al., 2025).

NGLY1-deficient patient-derived human induced pluripotent stem cell (hiPSC) and their neural derivatives provide an important human model for dissecting the mechanisms underlying the neurodevelopmental phenotypes of the disease. For example, cerebral organoid studies have shown that NGLY1 deficiency leads to premature neuronal differentiation and impairs the differentiation of SATB2+ cortical neurons, which are predominantly found in the upper layers of the cerebral cortex, potentially through dysregulation of cellular stress response pathways (Lin et al., 2022). Furthermore, reports on NGLY1 deficient midbrain organoids, neural progenitors, and neurons show protein aggregation, mitochondrial dysfunction, abnormal responses to oxidative stress, dysregulated autophagy, and synaptic abnormalities (Abbott et al., 2023; Manole et al., 2023; Shyr et al., 2025). However, these models have primarily focused on neurons, and the impact of NGLY1 deficiency on glial development and function remains largely unexplored.

Microglia are the resident immune cells of the central nervous system (CNS). They acquire their identity through developmental trajectories that differ from most circulating myeloid populations. Fate-mapping studies established that adult microglia arise from primitive macrophage progenitors (PMPs) rather than definitive hematopoiesis (Ginhoux et al., 2010). Molecular profiling further showed that mature microglial identity depends on specialized developmental and tissue cues (Butovsky et al., 2014). Human iPSC models now allow key features of microglia development to be studied in defined genetic backgrounds. Such systems have generated microglial cells with typical marker expression, inflammatory responsiveness, phagocytosis, and disease-relevant signaling properties (Abud et al., 2017; Guttikonda et al., 2021; Muffat et al., 2016; Xu et al., 2020). In addition, human microglia show species-specific transcriptional and functional features that are only partly reproduced in mouse models (Galatro et al., 2017; Gosselin et al., 2017; Jiang et al., 2020). Cross-species analyses further showed that human microglia display greater transcriptional heterogeneity than rodent microglia (Geirsdottir et al., 2019), and that disease-related myeloid programs in mouse models capture only part of the human neurodegenerative response (Friedman et al., 2018). These differences support the use of hiPSC-derived microglia as a complementary platform to study human-specific developmental and functional consequences of NGLY1 deficiency. Here, using patient-derived NGLY1-deficient hiPSCs and CRISPR-mediated, gene-corrected isogenic lines, we examined how loss of NGLY1 affects the development of microglial progenitors (i.e., PMPs) and microglia. Our findings suggest a previously unrecognized role for NGLY1 in microglia. Elucidating the glial contribution to the neurodevelopmental abnormalities caused by NGLY1 deficiency could generate novel mechanistic insights into the pathogenesis of NGLY1 deficiency and offer potential targets for therapeutic intervention.

## Results

### Generation of gene-corrected isogenic iPSC lines from NGLY1 deficiency patient-derived iPSCs

To determine whether NGLY1 deficiency affects human microglial development, we first performed gene correction in a patient derived hiPSC line NGLY1Pt1i-509, which was generated from dermal fibroblasts from an NGLY1-deficient patient. The patient carried two compound heterozygous NGLY1 variants, a 3-bp deletion in exon 8 (c.1205_1207delTTC; p.F402del) and a nonsense mutation in exon 11 (c.1570C>T; p.R542X) (Enns *et al*., 2014; Zolekar et al., 2018). The NGLY1 mutations were corrected by CRISPR-mediated gene editing and replaced with normal NGLY1 cDNA from exon 5 through exon 12 (**Figure 1A**). Sanger sequencing confirmed restoration of the normal NGLY1 cDNA sequence in correctly targeted hiPSC clones (**Figure 1B**), which were subsequently verified by PCR (**Supplementary Figure S1A**). We next examined whether genetic correction restored NGLY1 protein expression. Western blot analysis confirmed the absence of NGLY1 protein in NGLY1-deficient hiPSCs (**Figure 1C**). In contrast, gene correction restored NGLY1 protein expression in the isogenic repaired hiPSCs (**Figure 1C**). Both NGLY1-deficient and gene-corrected isogenic hiPSCs expressed OCT4, TRA1-81, and NANOG, and maintained the same karyotype (**Supplementary Figure S1B, C**). Taken together, gene correction restored NGLY1 expression while preserving pluripotent stem cell identity.

**Figure 1.**
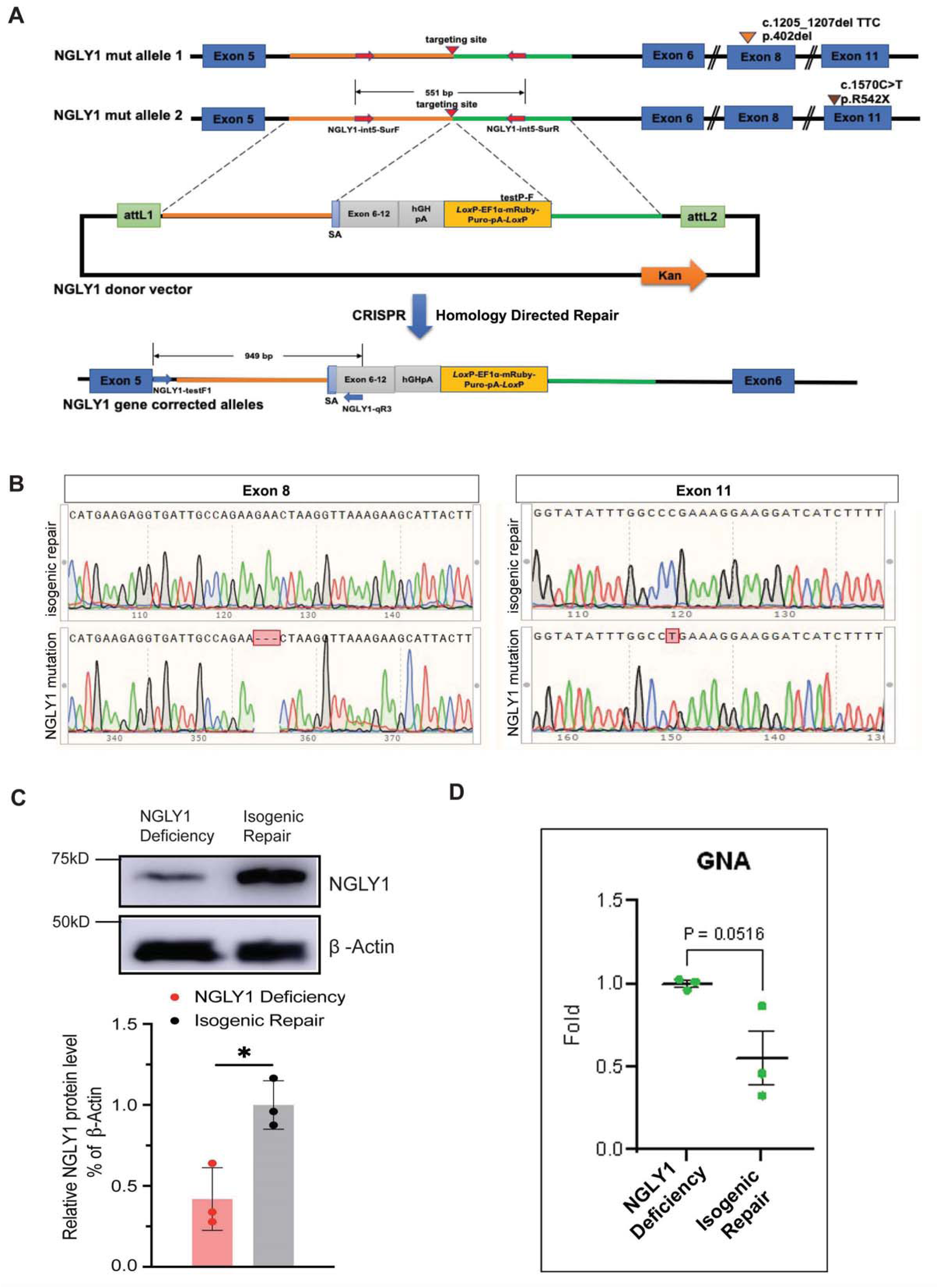
Generation and validation of CRISPR-mediated gene-corrected isogenic hiPSCs from NGLY1-deficient iPSCs. **(A)** Schematic of the CRISPR-mediated gene correction strategy in patient-derived NGLY1-deficient hiPSC. The maternally inherited NGLY1 mutation is located in exon 8, and the paternally inherited mutation is located in exon 11. The targeting vector was designed to replace exons 6 through 12 at both alleles simultaneously via CRISPR-mediated homology-directed repair. The sgRNA targeting sites, donor template, and expected correctly targeted allele structure following homology-directed repair are shown. The resulting gene-corrected hiPSC line restores the normal NGLY1 coding sequence. **(B)** Representative Sanger sequencing chromatogram showing validation of gene correction of NGLY1 cDNA at the corresponding sites in exon 8 and exon 11. Boxed regions highlight mutation sites. **(C)** Western blot analysis of NGLY1 protein expression in NGLY1-deficient and gene-corrected hiPSCs. NGLY1 expression was restored in gene-corrected hiPSCs but was barely detectable in NGLY1-deficient hiPSCs. Three independent experiments (n = 3) were performed. Data are presented as mean ± SEM. *p*<0.05 **(D)** Relative GNA levels. NGLY1-deficient PMPs have markedly higher GNA than gene-corrected isogenic repaired PMPs. Three independent experiments (n = 3) were performed. Data are presented as mean ± SEM. p=0.0516.

### NGLY1-deficient hiPSC-derived PMPs exhibit impaired proliferation

To test whether NGLY1 deficiency affects early microglial development, we differentiated the isogenic pair of patient and gene-corrected hiPSCs into PMPs in parallel, using a highly efficient and reproducible protocol (Haenseler et al., 2017; Jiang *et al*., 2020; Jin et al., 2026b; Xu et al., 2021) (**Figure 2A**). In both patient and gene-corrected isogenic groups, more than 90% of PMPs expressed CD235, a marker of yolk-sac primitive hematopoietic progenitors, and CD43, a hematopoietic progenitor marker (**Figure 2B, C**). Consistent with the result in hiPSCs, NGLY1 expression was only detected in gene-corrected PMPs but not in patient PMPs (**Supplementary Figure S2A**). We next examined whether NGLY1 deficiency affects the basic characteristics of PMPs. Elevated GNA level is a functional marker in NGLY1-deficiency. Impaired deglycosylation leads to the generation of GlcNAc-Asn (GNA) through alternative lysosomal degradation pathways, resulting in the accumulation of GNA in cells (Haijes et al., 2019; Mueller et al., 2022; Suzuki et al., 2016; Zhu et al., 2022). To confirm functional correction in the gene-corrected isogenic PMPs, we measured GNA levels by mass spectrometry. Compared with NGLY1-deficient PMPs, gene-corrected isogenic PMPs showed markedly reduced GNA levels (p = 0.051, **Figure 1D**).

**Figure 2.**
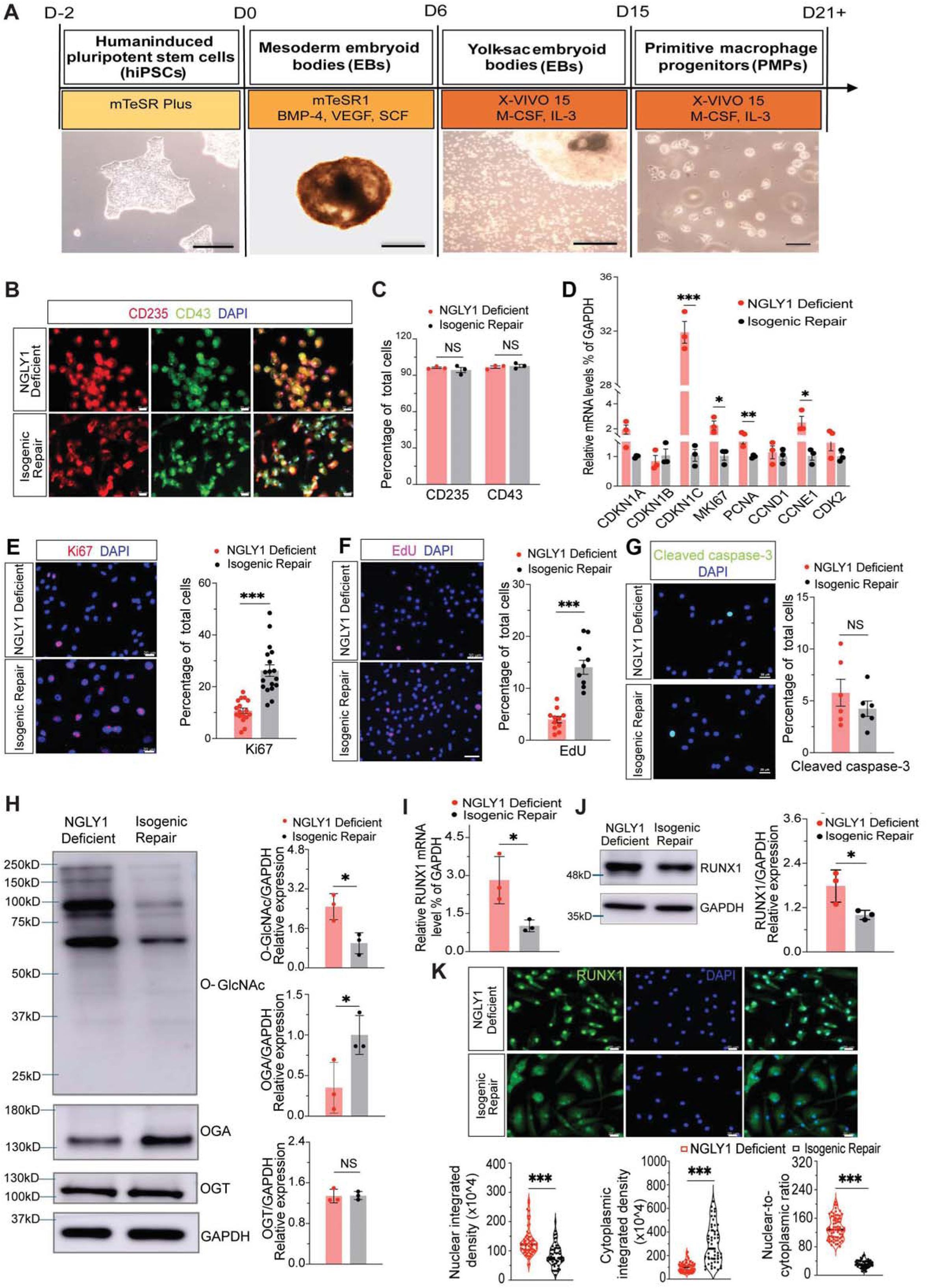
NGLY1 deficiency increases global O-GlcNAcylation and impairs proliferation in human iPSC derived PMPs. **(A)** Schematic overview of the differentiation protocol used to generate PMPs from hiPSCs. Scale bars, 100 μm in hiPSCs, mesoderm EBs, and yolk-sac EBs; 20 μm in PMPs. **(B)** Representative images of CD235 and CD43 immunocytochemistry staining in NGLY1-deficient and gene-corrected isogenic PMPs. Scale bars, 20 μm. **(C)** Quantification of CD235^+^ and CD43^+^ cells among total cells in NGLY1-deficient and gene-corrected PMPs. Similar percentages of CD235Ά and CD43Ά cells were detected in NGLY1-deficient and gene-corrected PMPs, indicating comparable PMP differentiation efficiency. Three independent experiments (n = 3) were performed. NS, not significant. Data are presented as mean ± SEM. **(D)** qPCR analysis of cell-cycle regulatory and proliferation-associated genes in PMPs. Transcript levels of CDKN1C, MKI67 (Ki-67), PCNA, and CCNE1 were significantly increased in NGLY1-deficient PMPs, while no significant differences were observed in CDKN1A, CDKN1B, CCND1, or CDK2. Three independent experiments (n = 3) were performed. \**p*<0.05, \*\**p*<0.01 and \*\*\**p*<0.001. Data are presented as mean ± SEM. **(E)** Representative images and quantification of Ki67^+^ cells in NGLY1-deficient and gene-corrected PMPs. The percentage of Ki-67+ cells was significantly lower in NGLY1-deficient PMPs. Each point represents one field of view. NGLY1-deficient, n =18 fields; Gene-corrected isogenic , n=19 fields. n = 3 independent differentiations per group; \*\*\**p*<0.001; data are presented as mean ± SEM. Scale bars, 20 μm. **(F)** Representative images and quantification of EdU-positive PMPs. EdU incorporation was significantly lower in NGLY1-deficient hiPSC-derived PMPs than in gene-corrected PMPs. Each point represents one field of view. NGLY1-deficient, n =11. Gene-corrected isogenic, n=9.; Fields are from n = 3 independent differentiations per group; \*\*\**p*<0.001; data are presented as mean ± SEM. Scale bars, 50 μm. **(G)** Representative images and quantification of cleaved caspase-3^+^ cells in PMPs by immunofluorescence staining. No significant differences were found in NGLY1-deficient and gene-corrected hiPSC derived PMPs. Each point represents one field of view. NGLY1-deficient, n =6; Gene-corrected isogenic, n=6. Fields from n = 3 independent differentiations per group; NS, not significant; data are presented as mean ± SEM. Scale bars, 20 μm. **(H)** Western blot analysis of global O-GlcNAcylation, OGA, and OGT in PMPs. Quantification revealed significantly higher O-GlcNAcylation levels in NGLY1-deficient PMPs. OGA protein levels were also significantly increased, while OGT protein expression was similar between the two groups. GAPDH was used as a loading control. Three independent experiments (n = 3) were performed. NS, not significant; \**p*<0.05. Data are presented as mean ± SEM. **(I)** qPCR analysis of *RUNX1* mRNA expression in PMPs. RUNX1 mRNA expression was significantly increased by more than twofold in NGLY1-deficient PMPs than in gene-corrected isogenic PMPs. Three independent experiments (n = 3) were performed. \**p*<0.05. Data are presented as mean ± SEM. **(J)** Western blot analysis of RUNX1 protein abundance in PMPs. RUNX1 protein expression was significantly increased by more than 1.5-fold in NGLY1-deficient PMPs than in gene-corrected isogenic PMPs. Three independent experiments (n = 3) were performed. \**p*<0.05. Data are presented as mean ± SEM. **(K)** Representative images and quantification of RUNX1 nuclear integrated density, cytoplasmic integrated density, and the nuclear-to-cytoplasmic ratio in PMPs. RUNX1 nuclear integrated density and the nuclear-to-cytoplasmic ratio were significantly increased, while cytoplasmic integrated density was significantly decreased in NGLY1-deficient PMPs than in gene-corrected isogenic PMPs, indicating increased nuclear translocation of RUNX1. Each point represents one field of view. NGLY1-deficient, n =55; Gene-corrected isogenic, n=51. Fields from n = 3 independent differentiations per group; \*\*\**p*<0.001; data are presented as mean ± SEM. Scale bars, 20 μm.

Immunofluorescence staining revealed a >15% reduction in the percentage of Ki-67+ cells in NGLY1-deficient PMPs compared with gene-corrected PMPs (p<0.001, **Figure 2E**). EdU incorporation was also reduced by approximately 10% in NGLY1-deficient PMPs (p<0.001, **Figure 2F**), indicating impaired cell proliferation.

Interestingly, mRNA expression of proliferation-associated genes, including MKI67, PCNA, and CCNE1, was increased in NGLY1-deficient PMPs as shown by quantitative PCR (qPCR) data (**Figure 2D)**. To further examine the effects of NGLY1 on additional cell cycle regulators, we performed qPCR analysis of CDKN1A, CDKN1B, and CDKN1C, which encode members of the Cip/Kip family of cyclin-dependent kinase inhibitors that restrain cell-cycle progression. While the mRNA expression of *CDKN1A* and *CDKN1B* remained unchanged, *CDKN1C* was significantly increased in NGLY1-deficient PMPs (p<0.001, **Figure 2D**). Consistent with the qPCR results, CDKN1C protein expression was also significantly increased (p<0.001, **Supplementary Figure S2B**). Meanwhile, cleaved Caspase-3 staining showed no significant difference, suggesting that NGLY1 deficiency had little or no effect on apoptosis (**Figure 2G**). Taken together, these results indicate that NGLY1 deficiency impairs cell-cycle progression and proliferation without substantially affecting apoptosis. The discrepancy between cell cycle-related gene expression and protein abundance suggests that post-transcriptional and/or post-translational regulatory mechanisms may contribute to altered cell-cycle progression in NGLY1 deficiency.

### NGLY1 deficiency disrupts O-GlcNAcylation homeostasis

NGLY1 and O-GlcNAc cycling are key regulators of glycoprotein homeostasis, affecting protein quality control, aggregation, and cell fate (Na et al., 2022; Sekine and Motohashi, 2021; Suzuki *et al*., 2016). O-GlcNAcylation also regulates transcriptional programs underlying essential cellular processes. We therefore examined whether O-GlcNAc cycling is altered in NGLY1-deficient cells and whether it contributes to the molecular dysregulation associated with NGLY1 deficiency. Western blot analysis using the RL2 antibody, which detects global O-GlcNAcylation, showed increased O-GlcNAc levels in NGLY1-deficient PMPs compared with gene-corrected PMPs (**Figure 2H**). This increase was accompanied by a 70% reduction in the protein expression of O-GlcNAcase (OGA), the enzyme responsible for removing O-GlcNAc modification from proteins. In contrast, no significant change was detected in the abundance of O-GlcNAc transferase (OGT), the enzyme responsible for adding an O-GlcNAc to target proteins (**Figure 2H**). Taken together, these data indicate that NGLY1-deficient PMPs exhibit elevated O-GlcNAcylation together with altered O-GlcNAc cycling enzyme balance.

### Transcription factor RUNX1 is upregulated in NGLY1 deficient PMPs

Because transcriptional networks govern cell fate and O-GlcNAc cycling regulates transcriptional programs, we next asked whether disruption of O-GlcNAc homeostasis in NGLY1-deficient cells alters key developmental transcriptional regulators involved in microglial lineage differentiation such as RUNX1, a transcription factor critical for hematopoietic emergence and early myeloid progenitor formation (Lancrin et al., 2009). Both RUNX1 transcript and protein levels were significantly increased in NGLY1-deficient PMPs as validated by qPCR and Western blot analysis (p<0.05 for both, **Figure 2I, J**). Furthermore, immunofluorescence analysis revealed increased nuclear RUNX1 integrated density and an elevated nuclear-to-cytoplasmic RUNX1 ratio in NGLY1-deficient PMPs (**Figure 2K**), indicating enhanced RUNX1 transcriptional activity. Together, these findings demonstrate that at the PMP stage, NGLY1 deficiency is associated with elevated O-GlcNAcylation, increased RUNX1 expression, and enhanced nuclear localization of RUNX1, likely driving aberrant activation of the microglial developmental program.

O-GlcNAcylation is a key regulator of transcriptional programs. To test whether O-GlcNAc cycling was linked to changes of RUNX1 in NGLY1 deficient PMPs, we pharmacologically modulated O-GlcNAcylation by OGA inhibitor Thiamet-G and OGT inhibitor OSMI-1. As expected, dose-response experiments showed that 24 h treatment with the OGA inhibitor Thiamet-G increased global O-GlcNAcylation in gene-corrected PMPs, while the OGT inhibitor OSMI-1 decreased O-GlcNAcylation in NGLY1-deficient PMPs (**Supplementary Figure S2C, D**). Based on these dose responses, we selected 5 μM Thiamet-G and 10 μM OSMI-1 for subsequent experiments. In gene-corrected PMPs, Thiamet-G treatment inhibited OGA, increased O-GlcNAcylation, and increased RUNX1 protein abundance. Conversely, in NGLY1-deficient PMPs, OSMI-1 treatment inhibited OGT, leading to reduced O-GlcNAcylation and a corresponding decrease in RUNX1 protein level (**Supplementary Figure S2E-G**). Together, our findings suggest that altered O-GlcNAc cycling may contribute to RUNX1 dysregulation in NGLY1-deficient PMPs.

### Differential transcriptomic profiles between NGLY1-deficient and gene-corrected PMPs

To determine global transcriptional changes associated with NGLY1 deficiency, we performed bulk RNA sequencing (RNA-Seq) on NGLY1-deficient PMPs and their gene-corrected isogenic counterparts. As shown in **Figure 3A**, expression of 2,874 genes was significantly upregulated, and 1,323 genes downregulated in NGLY1-deficient PMPs. Among the differentially expressed genes (DEGs) that were significantly decreased, the immune and myeloid identity genes stood out, which included *HLA-DRA, HLA-DRB1, HLA-DQA1, HLA-DPA1, CIITA, SPI1, TREM2, TYROBP,* and *CX3CR1* (**Figure 3A**). In contrast, expression of developmental and remodeling-associated genes, including *COL3A1, FN1, and CTGF*, was increased. To gain deeper insight into the biological processes and molecular pathways altered in these cells, we performed gene ontology (GO) enrichment analysis and gene set enrichment analysis (GSEA) of the DEGs identified from the comparison of NGLY1-deficient and gene-corrected PMPs. GO analysis showed that DEGs downregulated in NGLY1-deficient PMPs were enriched for immune, antigen-presentation, leukocyte, and myeloid lineages (**Figure 3B, Supplementary Figures S3A, C**), which was confirmed by qPCR showing decreased expression of selected immune and myeloid genes such as *HLA-DRA, HLA-DRB1, TYROBP, TREM2,* and *SPI1* (**Figure 3D**). Further analysis of the downregulated DEGs by GSEA revealed significant downregulation of the N-glycan trimming/calnexin-calreticulin cycle and G1/S DNA damage checkpoint pathways in NGLY1-deficient PMPs (**Supplementary Figures S3B-E**), indicating impaired ER protein quality control and altered cell-cycle checkpoint regulation following NGLY1 deficiency. In contrast, DEGs upregulated in NGLY1-deficient PMPs were associated with extracellular matrix organization, cell adhesion, and neurodevelopmental remodeling pathways (**Figure 3C**). Consistent with these transcriptomic findings, qPCR confirmed increased expression of COL3A1, FN1, and CTGF in NGLY1 deficient PMPs (**Figure 3D**).

**Figure 3.**
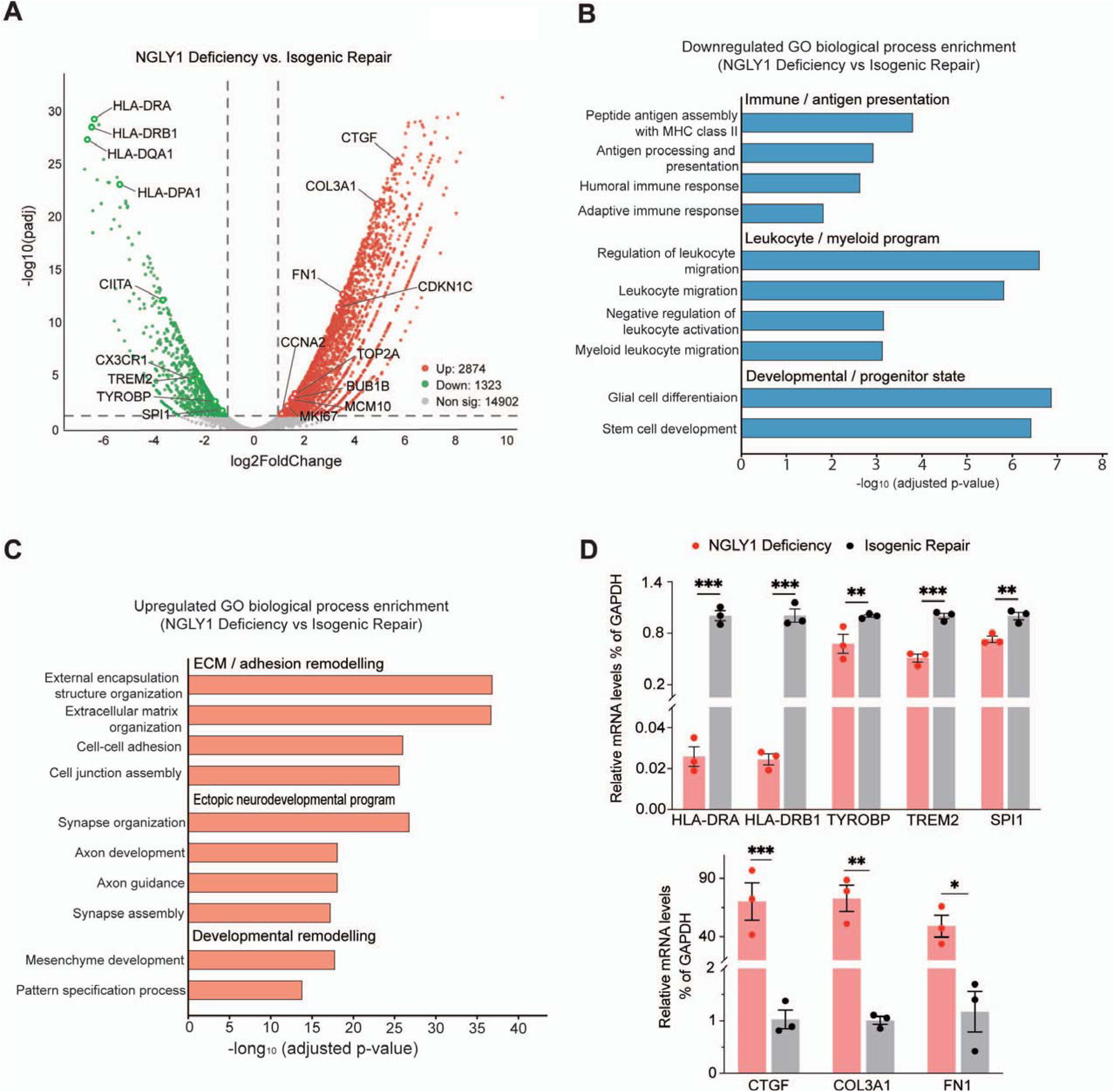
Transcriptomic profiling of NGLY1-deficient PMPs vs. gene-corrected isogenic, PMPs. **(A)** Volcano plot showing RNA-Seq results comparing NGLY1-deficient and gene-corrected PMPs. Relative to gene-corrected PMPs, differentially expressed genes (DEGs) that were downregulated in NGLY1-deficient PMPs are shown in green, and upregulated DEGs are shown in red. **(B)** Gene Ontology biological process enrichment analysis of top 10 downregulated terms in NGLY1-deficient vs. gene-corrected PMPs. **(C)** Gene Ontology biological process enrichment analysis of top 10 upregulated terms in NGLY1-deficient vs. gene-corrected PMPs. **(D)** qPCR validation of gene expression of selected DEGs identified by RNA-Seq in NGLY1-deficient and gene-corrected PMPs. The upper panel shows immune/myeloid-associated genes, including HLA-DRA, HLA-DRB1, TYROBP, TREM2, and SPI1. The lower panel shows developmental/remodeling-associated genes, including CTGF, COL3A1, and FN1. Three independent experiments (n = 3) were performed for each assay. \**p*<0.05. Data are presented as mean ± SEM.

### NGLY1-deficient PMPs produce microglia with sustained functional abnormalities

To test whether the abnormalities observed in NGLY1-deficient PMPs persisted following microglial differentiation, PMPs derived from patient NGLY1-deficient hiPSCs and their gene-corrected isogenic cells were further differentiated into microglia. (**Figure 4A**). Both lines generated microglia expressing canonical microglial markers IBA1, TMEM119, and CX3CR1 at similar efficiencies in vitro in culture (**Figure 4B**). To assess the basic functions of NGLY1-deficient microglia, we next examined their responses to inflammatory stimuli. Under unstimulated conditions, inflammatory response genes *IL1B*, *IL6*, and *TNF (TNFα)* were expressed at higher levels in NGLY1-deficient than in gene-corrected microglia, as shown in qPCR analysis (**Figure 4C**). Upon LPS stimulation, the increase of gene expression for *IL1B*, *IL6*, and *TNF* was even more pronounced in NGLY1 deficient microglia, especially for genes *IL1B* and *IL6*. To determine whether elevated expression of inflammatory cytokines was accompanied by microglial activation, we examined CD68, a lysosomal protein commonly used as a marker of microglial activation and phagocytic activity. As expected, both NGLY1-deficient and gene-corrected microglia showed elevated CD68 expression following LPS stimulation. However, although LPS increased CD68 expression in both groups, the magnitude of this increase was significantly attenuated in NGLY1-deficient microglia compared with gene-corrected microglia (**Figures 4D, E**), indicating an impaired activation response to LPS in NGLY1-deficient microglia. Upon activation by inflammatory or injury-related stimuli, microglia typically shift from a ramified morphology, characterized by elongated and highly branched processes, toward a more amoeboid morphology with shortened processes and a rounded cell body (Prinz et al., 2019). To assess changes of microglial morphology of NGLY1-deficient microglia upon LPS exposure, we quantified the outlines of IBA1^+^ microglia using FracLac in ImageJ (**Figure 4F**). Compared with gene-corrected microglia, NGLY1-deficient microglia showed lower fractal dimension, higher circularity, and reduced spine ratio under basal and LPS-stimulated conditions (**Figure 4G**). These results indicate that NGLY1-deficient microglia exhibit decreased morphological complexity in response to LPS exposure. Microglial phagocytosis is required for apoptotic cell clearance, synaptic refinement, and immune homeostasis (Paolicelli et al., 2011; Schafer et al., 2012; Sierra et al., 2010). To test whether the impaired elevation of CD68 expression in response to LPS was associated with reduced phagocytic function, we measured functional phagocytosis using pH-sensitive pHrodo Zymosan beads (Hubschmann et al., 2022; Padmakumar et al., 2023). As expected, NGLY1-deficient microglia showed reduced phagocytic capacity compared with gene-corrected microglia across all LPS concentrations tested (**Figure 4H, I**), confirming that NGLY1-deficient microglia have dysregulated inflammatory gene expression, reduced microglial activation, and impaired phagocytic capacity.

**Figure 4.**
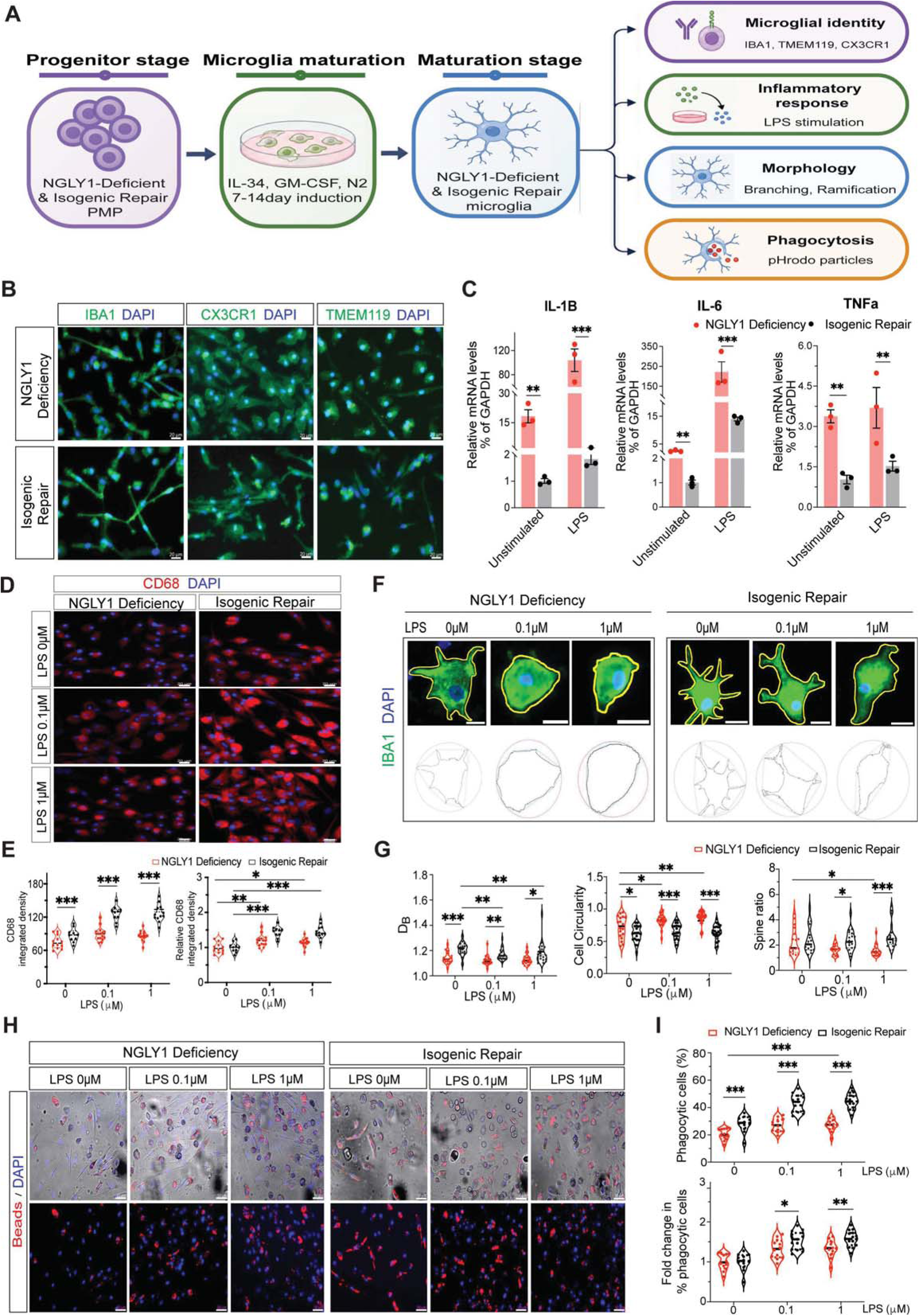
Impaired glycoprotein degradation produces persistent microglial dysfunction in hiPSC-derived microglia. **(A)** A schematic diagram of the experimental workflow, created using BioRender.com. **(B)** Representative images of hiPSC-derived microglia stained for IBA1, CX3CR1, and TMEM119. No significant differences in differentiation efficiency, percentages of marker-positive cells, or cell morphology were observed between NGLY1-deficient and gene-corrected hiPSC-derived microglia. Scale bars, 20 μm. **(C)** qPCR analysis of *IL1B*, *IL6*, and *TNFα* gene expression in NGLY1-deficient and gene-corrected hiPSC-derived microglia under unstimulated conditions or following LPS stimulation (0.1 μM) for 24 h. *IL1B*, *IL6*, and *TNFα* gene expression was significantly elevated under both unstimulated and LPS-stimulated conditions in NGLY1-deficient microglia. Three independent experiments (n = 3) were performed for each assay. \*\**p*<0.01, \*\*\**p*<0.001. Data are presented as mean ± SEM. **(D)** Representative images of CD68 immunofluorescence staining of microglia derived from NGLY1-deficient and gene-corrected hiPSCs following 24 h of LPS exposure at 0 μM, 0.1 μM or 1 μM. Scale bars, 20 μm. **(E)** Quantification of CD68 expression level. Fluorescence integrated density and Relative integrated density (normalized to each 0 μM LPS replicate) are shown. Following LPS treatment, CD68 integrated density and relative integrated density both were significantly lower in NGLY1-deficient microglia than in gene-corrected cells. Each point represents one field of view. NGLY1-deficient: 0 μM LPS n =12, 0.1 μM LPS n =15, 1 μM LPS n =14; Gene-corrected isogenic: 0 μM LPS n =10, 0.1 μM LPS n =10, 1 μM LPS n =12. Fields from n = 3 independent differentiations per genotype. \**p*<0.05, \*\**p*<0.01 and \*\*\**p*<0.001. Data are presented as mean ± SEM. **(F)** Representative IBA1 immunofluorescence staining images showing regions of interest (ROI)-based tracing of NGLY1-deficient and gene-corrected isogenic hiPSC-derived microglia from treated with 0, 0.1, or 1 μM LPS. The upper panels show IBA1^+^ microglia used for morphological analysis. Lower panels show the corresponding binary outlines and trace cellular regions used to quantify fractal dimension, cell circularity, and spine ratio. NGLY1-deficient microglia show less ramified morphology and increased circularity. Scale bars, 20 μm. **(G)** Morphological analysis of IBA1^+^ microglia using the FracLac plugin in ImageJ. Representative images of IBA1^+^ microglia were analyzed using the FracLac plugin in ImageJ to quantify morphological features. Quantitative analysis of fractal dimension (D_B_), circularity, and spine ratio revealed significant morphological differences between NGLY1-deficient and gene-corrected isogenic microglia. Compared with gene-corrected microglia, NGLY1-deficient microglia exhibited a significantly lower fractal dimension (D_B_), indicating reduced morphological complexity and branching, a significantly higher circularity, consistent with a more rounded morphology, and a significantly lower spine ratio, indicating fewer cellular protrusions. These findings show impaired microglial maturation and reduced ramification in NGLY1-deficient microglia. Analysis was performed on n = 18, 20, and 20 cells from NGLY1-deficient microglia and n = 18, 20, and 20 cells from gene-corrected microglia following treatment with 0, 0.1, and 1 μM LPS, respectively, across 3 independent differentiations per group. \**p*<0.05, \*\**p*<0.01 and \*\*\**p*<0.001. Data are presented as mean ± SEM. **(H)** Representative images of pHrodo particle uptake by NGLY1-deficient and gene-corrected microglia stimulated with 0, 0.1, or 1 μM LPS. The top panels show red fluorescent beads internalized by microglia, together with nuclear counter stain (DAPI) and bright-field images. The bottom panels show red fluorescent beads within phagocytic microglia together with DAPI from the corresponding fields. Under both unstimulated (0 μM LPS) and LPS-stimulated conditions (0.1 and 1 μM LPS), NGLY1-deficient microglia exhibited significantly reduced phagocytic activity compared with gene-corrected isogenic microglia. Scale bars, 50 μm. **(I)** Quantification of the percentage of phagocytic cells and relative fold change of the percentage of phagocytic cells normalized to each 0 μM LPS replicate. NGLY1-deficient microglia show decreased percentage of phagocytic cells and relative fold change. Phagocytosis analysis was performed on n = 12, 12, and 12 fields of view from NGLY1-deficient microglia and n = 12, 15, and 15 fields of view from gene-corrected microglia for 0, 0.1, and 1 μM LPS, respectively, across 3 independent differentiations per genotype. \**p*<0.05, \*\**p*<0.01 and \*\*\**p*<0.001. Data are presented as mean ± SEM.

### RUNX1 knockdown attenuates selected phenotypes of NGLY1-deficient PMP and microglia

Gene expression of RUNX1 was increased in NGLY1-deficient PMPs. Given the critical role of RUNX1 in myeloid lineage development and differentiation, we next tested whether perturbation of RUNX1 expression could restore normal PMP differentiation and functional phenotypes in NGLY1 deficient cells. We first established stable RUNX1 knockdown in NGLY1-deficient hiPSCs using a RUNX1-targeting shRNA (shRNA749 or shRNA750), with a non-targeting (NT) shRNA as the control (**Figure 5A, Supplementary Figure S4A**). qPCR results showed RUNX1 expression was reduced by approximately 50% compared to gene-corrected isogenic repaired line, and was reduced by approximately 83% to NT shRNA control, confirming efficient RUNX1 knockdown in PMPs (**Figure 5B**). Western blot further confirmed a 70% reduction in RUNX1 protein levels in PMPs expressing either RUNX1 shRNA749 or shRNA750 (**Figure 5C, D**). At the undifferentiated stage, RUNX1 shRNA-expressing hiPSCs retained OCT4 and TRA1-81 expression. Upon induction, these cells were able to generate CD43^+^ and CD235^+^ PMPs and IBA1^+^ microglia at a similar efficiency to the parental NGLY1-deficient hiPSCs (**Supplementary Figure S4B-D**). We then examined whether RUNX1 knockdown rescued the proliferation defects in NGLY1-deficient PMPs. Compared with the NT shRNA control, RUNX1 knockdown in NGLY1-deficient PMPs increased the proportion of Ki67^+^ and EdU^+^ cells by 15%, as shown by immunofluorescence staining experiments (**Figure 5E-H**). Consistently, the expression of *CDKN1C*, a key inhibitor of cell cycle progression, was also reduced in PMPs with RUNX1 knockdown, further supporting that RUNX1 knockdown at least partially restored the proliferative capacity of NGLY1-deficient PMPs (**Figure 5I**).

**Fig 5.**
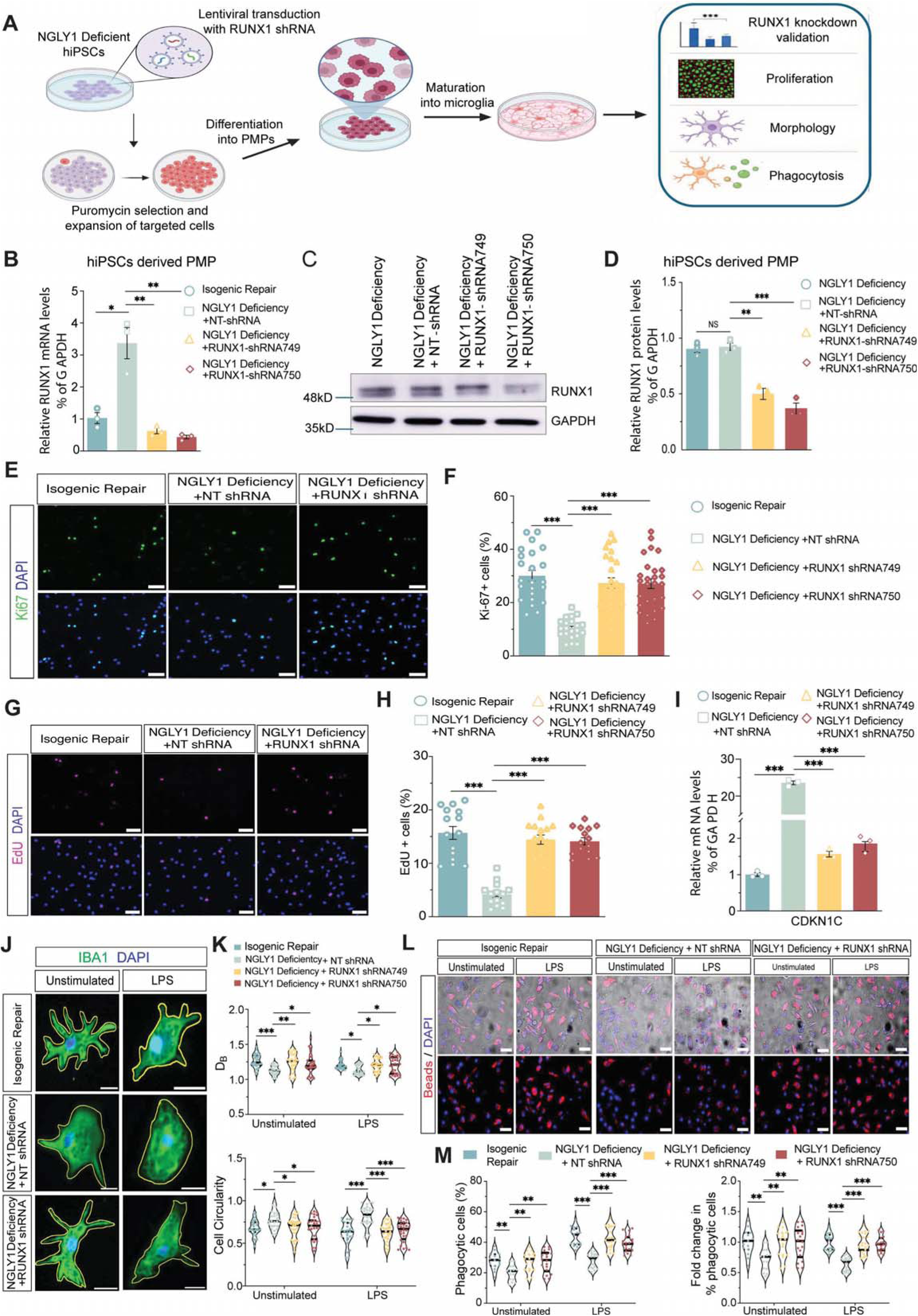
Perturbation of RUNX1 gene expression by shRNA partially restores phenotypes in NGLY1 deficient PMPs. **(A)** A schematic diagram showing the design of the *RUNX1* knockdown experiment. This drawing was created using BioRender.com. **(B)** qPCR analysis of *RUNX1* mRNA expression in hiPSC-derived PMPs expressing non-targeting (NT) shRNA, RUNX1 shRNA749, or RUNX1 shRNA750. Compared to untreated group, the RUNX1 mRNA expression was reduced by approximately 50% in shRNA 749 or shRNA 750 groups. Compared to NT shRNA, the RUNX1 mRNA expression was reduced by approximately 85% in shRNA 749 or shRNA 750 groups. Three independent experiments (n = 3) were performed. \**p*<0.05, \*\**p*<0.01. Data are presented as mean ± SEM. **(C)** Western blot analysis of RUNX1 protein abundance in hiPSC-derived PMPs expressing non-targeting (NT) shRNA, RUNX1 shRNA749, or RUNX1 shRNA750. **(D)** Quantification of RUNX1 protein expression shows that compared to untreated or NT shRNA groups, RUNX1 protein expression was reduced by approximately 50% in shRNA 749 or shRNA 750 groups. Three independent experiments (n = 3) were performed. \*\**p*<0.01, \*\*\**p*<0.001. Data are presented as mean ± SEM. **(E)** Representative images of Ki67 immunofluorescence staining in PMPs derived from gene-corrected isogenic hiPSCs, NGLY1-deficient hiPSCs expressing NT shRNA, and NGLY1-deficient hiPSCs expressing RUNX1 shRNA. Scale bars, 100 μm. **(F)** Quantification of the percentage of Ki67^+^ cells in PMPs in gene-corrected, NGLY1-deficient + NT shRNA, and NGLY1-deficient + RUNX1 shRNA groups. The percentage of Ki-67Ά cells was significantly lower in NGLY1-deficient PMPs than in gene-corrected PMPs. In contrast, shRNA-mediated RUNX1 knockdown partially restored the percentage of Ki-67Ά cells to levels comparable to those in the gene-corrected group. Each point represents one field of view; gene-corrected, n=23; NGLY1-deficient+ NT shRNA, n =21; NGLY1-deficient + RUNX1 shRNA749, n =26; NGLY1-deficient + RUNX1 shRNA750, n =27. Fields from N = 3 independent differentiations per genotype. \*\*\**p*<0.001. Data are presented as mean ± SEM **(G)** Representative images of EdU incorporation in PMPs in gene-corrected, NGLY1-deficient + NT shRNA, and NGLY1-deficient + RUNX1 shRNA groups. Scale bars, 100 μm. **(H)** Quantification of EdU^+^ cells in PMPs in gene-corrected, NGLY1-deficient + NT shRNA, and NGLY1-deficient + RUNX1 shRNA groups. The percentage of EdUΆ cells was significantly lower in NGLY1-deficient PMPs than in gene-corrected PMPs. In contrast, shRNA-mediated RUNX1 knockdown partially restored the EdUΆ incorporation to levels comparable to those in the gene-corrected group. Each point represents one field of view; gene-corrected, n=15; NGLY1-deficient+ NT shRNA, n =14; NGLY1-deficient + RUNX1 shRNA749, n =16; NGLY1-deficient + RUNX1 shRNA750, n =15. Fields from N = 3 independent differentiations per genotype. \*\*\**p*<0.001. Data are presented as mean ± SEM **(I)** qPCR analysis of *CDKN1C* expression in PMPs in gene-corrected, NGLY1-deficient + NT shRNA, and NGLY1-deficient + RUNX1 shRNA groups. Although CDKN1C mRNA expression was significantly elevated in the NT shRNA group, consistent with the NGLY1-deficient phenotype, CDKN1C expression in the shRNA749-and shRNA750-treated RUNX1 knockdown PMPs was restored to levels comparable to those in the gene-corrected group. Three independent experiments (n = 3) were performed for each experiment. \**p*<0.05, \*\**p*<0.01. Data are presented as mean ± SEM. **(J)** Representative IBA1 immunofluorescence staining images showing regions of interest (ROI)-based tracing of iPSC-derived microglia in gene-corrected, NGLY1-deficient + NT shRNA, and NGLY1-deficient + RUNX1 shRNA groups under unstimulated and LPS treated conditions. Compared with the NT shRNA group, microglia in the RUNX1 shRNA groups exhibited a more ramified morphology and reduced circularity, restoring their morphology toward that of the gene-corrected group. Scale bars, 20 μm. **(K)** Morphological analysis of microglia using the FracLac plugin in ImageJ. Compared with the NT shRNA group, RUNX1 shRNA-treated microglia exhibited restored fractal dimension (Db), reduced circularity, and an increased spine ratio to levels comparable to those of the gene-corrected group, indicating partial restoration of microglial maturation. Analysis was performed on n = 25 and 26 cells from gene-corrected microglia; n = 24 and 25 cells from NGLY1-deficient + NT shRNA microglia; n = 28 and 28 cells from NGLY1-deficient + RUNX1 shRNA749 microglia; n = 27 and 28 cells from NGLY1-deficient + RUNX1 shRNA750 microglia under unstimulated and LPS conditions, respectively, across 3 independent differentiations per genotype. \**p*<0.05, \*\**p*<0.01 and \*\*\**p*<0.001. Data are presented as mean ± SEM. **(L)** Representative images of pHrodo particles uptake in hiPSC-derived microglia in gene-corrected, NGLY1-deficient + NT shRNA, and NGLY1-deficient + RUNX1 shRNA groups under unstimulated and LPS-stimulated conditions. The upper images are overlapped with bright-field images for reference. Scale bars, 50 μm. **(M)** Quantification of the percentages of phagocytic cells and relative fold change of the percentage of phagocytic cells normalized to each 0 μM LPS replicate. Compared with the NT shRNA group, the RUNX1 shRNA knockdown groups exhibited partially restored phagocytic activity toward the levels observed in the gene-corrected microglia. Phagocytosis analysis was performed on n = 15 and 17 fields of view from gene-corrected microglia; n = 15 and 16 fields of view from NGLY1-deficient + NT shRNA microglia; n = 17 and 19 fields of view from NGLY1-deficient + RUNX1 shRNA749 microglia; n = 17 and 19 fields of view from NGLY1-deficient + RUNX1 shRNA750 microglia under unstimulated and LPS treated conditions, respectively, across 3 independent differentiations per genotype. \**p*<0.05, \*\**p*<0.01 and \*\*\**p*<0.001. Data are presented as mean ± SEM.

We next asked whether stable RUNX1 perturbation in hiPSCs affected downstream microglial phenotypes. Microglia derived from RUNX1-knockdown NGLY1-deficient hiPSCs showed a partial restoration of morphology, characterized by significant changes in fractal dimension and circularity (**Figure 5J, K**). The morphology more closely resembled that of microglia derived from gene-corrected hiPSCs (**Figure 4F**). In addition, phagocytosis assays using pHrodo Zymosan beads uptake demonstrated that RUNX1 knockdown partially restored the phagocytic capacity of NGLY1-deficient hiPSC-derived microglia to levels comparable to those of gene-corrected microglia under both unstimulated and LPS-stimulated conditions (**Figure 5L, M**).

Together, these findings indicate that RUNX1 knockdown initiated at the hiPSC stage partially rescues the defects of PMP proliferation and microglial differentiation and function in NGLY1-deficient cells.

## Discussion

Using NGLY1-deficient patient-derived hiPSCs and their corresponding gene-corrected isogenic cells, we found that NGLY1 deficiency disrupts microglial lineage differentiation and function, with abnormalities emerging at the PMP stage. These findings suggest that altered progenitor development may contribute to subsequent microglial dysfunction. Beyond GNA accumulation, increased global O-GlcNAcylation accompanied by reduced OGA expression and increased RUNX1 expression and nuclear localization suggests a potential link between disrupted O-GlcNAc regulation and altered myeloid development. Partial rescue of the cellular defects by RUNX1 knockdown supports RUNX1 dysregulation as a contributing mechanism.

NGLY1 plays a critical role in protein deglycosylation and glycoprotein quality control. Previous studies have primarily focused on its role in protein quality control and NRF1/NFE2L1-mediated proteasome recovery and cellular stress adaptation. For example, NGLY1-dependent peptide editing of NRF1 has been shown to regulate proteasome gene expression in multiple model systems. Accordingly, pharmacological or genetic inhibition of NGLY1 disrupts NRF1 processing, thereby impairing the proteasome recovery response. Loss of function of NGLY1 has also been linked to mitochondrial and immune dysregulation through the NGLY1-NRF1 axis (Lehrbach *et al*., 2019; Pandey et al., 2022; Tomlin *et al*., 2017; Yang *et al*., 2018). Neural differentiation of patient derived hiPSCs further show that NGLY1 deficiency can impair neuronal protein homeostasis and neural development (Abbott *et al*., 2023; Lin *et al*., 2022; Manole *et al*., 2023).

Aligned with observations in the neural lineage, our data identify the PMP stage as an early cellular target of NGLY1 deficiency, extending the impact of NGLY1 beyond neural progenitors and neurons to the developing microglia. Specifically, our results in NGLY1-deficient PMPs and microglia establish a previously underappreciated link between protein deglycosylation and O-GlcNAc cycling. Loss of NGLY1 function disrupts protein deglycosylation and, in parallel, increases global O-GlcNAcylation through downregulation of O-GlcNAcase (OGA), the enzyme responsible for removing O-GlcNAc modifications from proteins. Elevated O-GlcNAc signaling, in turn, increases the protein abundance of RUNX1, thereby decreasing cell proliferation while promoting premature PMP differentiation and impairing subsequent microglial maturation. Collectively, these data suggest that NGLY1 deficiency alters the myeloid progenitor state, leading to subsequent microglial dysfunction.

O-GlcNAcylation is a dynamic, nutrient- and stress-responsive post-translational protein modification in which O-GlcNAc transferase (OGT) adds, and O-GlcNAcase (OGA) removes, O-GlcNAc from serine and threonine residues of intracellular proteins (Yang and Qian, 2017). O-GlcNAcylation has emerged as a critical post-translational modification that regulates diverse biological processes by modulating the activity of transcription factors, thereby influencing their stability, subcellular localization, DNA binding, protein-protein interactions, and transcriptional activity (Hart et al., 2011; Parker et al., 2021; Yang and Qian, 2017). For example, O-GlcNAcylation enhances NF-κB-dependent inflammatory gene expression (Yang et al., 2008), increases the stability and transactivation activity of Sp1 (Han and Kudlow, 1997; Yang et al., 2001), promotes FOXO1 nuclear localization and transcriptional activity (Housley et al., 2008), antagonizes phosphorylation-dependent degradation of c-Myc (Chou et al., 1995), and stabilizes p53 by inhibiting MDM2-mediated degradation (Yang et al., 2006). Collectively, these studies demonstrate that O-GlcNAcylation serves as a central regulatory mechanism linking cellular metabolic status to transcriptional programs that control proliferation, lineage commitment, stress responses, and inflammation. A recent report (Zhou et al., 2025) demonstrated that RUNX1 is a direct target of O-GlcNAcylation. The authors showed that RUNX1 physically interacts with OGT and is directly modified by O-GlcNAcylation. Knockdown of OGT reduced RUNX1 O-GlcNAcylation and protein abundance without affecting RUNX1 mRNA levels, while OGT overexpression increased RUNX1 protein expression, indicating that O-GlcNAcylation regulates RUNX1 at the post-translational level and that O-GlcNAcylation stabilizes RUNX1 by preventing ubiquitination and subsequent protein degradation. Another study reported that RUNX1 functions downstream of O-GlcNAc signaling and mediates the effects of high glucose-induced O-GlcNAcylation on endothelial cell proliferation and migration (Xing et al., 2021). Our data are consistent with and extend these previous reports. NGLY1-deficient PMPs exhibited reduced OGA expression, increased global O-GlcNAcylation, and elevated RUNX1 protein levels. Moreover, pharmacological manipulation of O-GlcNAc cycling further supported this relationship: inhibition of OGA with Thiamet-G increased RUNX1 expression, while inhibition of OGT with OSMI-1 reduced RUNX1 levels. Because O-GlcNAc homeostasis is maintained by the opposing activities of OGT and OGA, our data suggest that reduced OGA expression is sufficient to promote RUNX1 accumulation. Whether OGA directly regulates RUNX1 remains unknown. Given its role as the enzyme that removes O-GlcNAc modifications, reduced OGA expression could lead to increased O-GlcNAcylation and stability of RUNX1 without requiring a direct OGA-RUNX1 interaction. This possibility warrants further investigation. Together, these findings suggest that elevated O-GlcNAcylation promotes RUNX1 accumulation in NGLY1-deficient PMPs, providing a mechanistic link between disrupted protein deglycosylation, altered O-GlcNAc homeostasis, and abnormal myeloid lineage commitment.

Interestingly, our data show that NGLY1-deficient PMPs exhibit impaired cell proliferation. Immunocytochemical analysis revealed a 2.5-fold reduction in Ki-67 protein expression, which was further supported by decreased EdU incorporation, the gold standard for measuring cell proliferation (Salic and Mitchison, 2008). These results are corroborated by increased expression of CDKN1C (p57^KIP2^), a member of the Cip/Kip family of cyclin-dependent kinase (CDK) inhibitors that regulates stem cell proliferation and differentiation, cell-cycle exit, hematopoietic development and maturation (Borges et al., 2015; Matsuoka et al., 1995). Importantly, several cell cycle- and proliferation-associated genes exhibited a discrepancy between mRNA and protein expression. For example, MKI67 mRNA levels were increased approximately threefold in NGLY1-deficient PMPs compared with gene-corrected isogenic PMPs, as determined by both qPCR and RNA-seq, despite a marked reduction in Ki-67 protein expression. This discrepancy may reflect post-transcriptional regulatory mechanisms, altered protein translation or stability, and/or compensatory transcriptional feedback in response to reduced protein abundance.

Previous reports in NGLY1-deficient animal or cell models have reported little or no obvious defects in the glial lineage, although microglia were not specifically or comprehensively examined (Asahina *et al*., 2021a; Manole *et al*., 2023; Yang *et al*., 2025). Using patient-derived isogenic hiPSC models, we showed that NGLY1 deficiency causes marked defects in both microglial development and maturation. These abnormalities originate at the PMP stage, where impaired proliferation and altered developmental programs precede defective microglial maturation and function. Together, these findings uncover a previously unrecognized role for NGLY1 in regulating human microglial lineage development. In summary, our results support a model in which loss of NGLY1 reprograms the early PMP state, leading to downstream defects in microglial maturation and function.

### Limitations of the study

This study has several limitations. First, although our results show that NGLY1 deficiency increases global O-GlcNAcylation, elevates RUNX1 abundance and nuclear localization, and that RUNX1 knockdown partially rescues the observed phenotypes, the precise molecular mechanisms linking NGLY1 deficiency to altered O-GlcNAc cycling and RUNX1 regulation remain to be defined. In particular, how NGLY1 deficiency leads to reduced OGA expression and increased global O-GlcNAcylation, whether OGA directly or indirectly regulates RUNX1 stability, and whether OGA physically interacts with RUNX1 require further investigation. Elucidating these molecular mechanisms will provide important insight into how dysregulated O-GlcNAc cycling influences microglial lineage specification Second, our conclusions are based on in vitro differentiation of patient-derived hiPSCs into PMPs and microglia. Although this platform recapitulates key stages of human microglial development, it does not fully capture the complex environment of the developing brain. Future studies assessing the differentiation, maturation, and function of NGLY1-deficient PMPs and microglia following transplantation into the developing brain, together with analyses of microglial phenotypes in NGLY1-deficient mice, would help determine whether the defects observed in vitro persist in vivo and contribute to altered brain development and neurological dysfunction. Finally, although RUNX1 knockdown partially rescued defects in PMP proliferation, microglial morphology, and phagocytic function, the rescue was incomplete, suggesting that additional pathways downstream of NGLY1 deficiency contribute to microglial dysfunction. Identifying these complementary mechanisms will further improve our understanding of the molecular basis of NGLY1 deficiency and may reveal additional therapeutic targets.

## Materials and Methods

### Vector construction for gene correction

For patient 509 hiPSCs, patient fibroblasts were obtained from the Coriell Institute for Medical Research. Whole genome sequencing showed that the patient is compound heterozygous for two NGLY1 mutations. The mutation in the maternal allele is a 3-base pair deletion in exon 8 (c.1205_1207del), in which TTC was deleted, resulting in an in-frame deletion of phenylalanine at position 402 (p.F402del; chromosome 3:25775416). The mutation in the paternal allele is a nonsense mutation in exon 11 (c.1570C>T), causing a change from arginine to a stop codon at amino acid position 542 (p.R542X; chromosome 3:25761670). To perform gene correction for both alleles simultaneously, human NGLY1 (exons 6 to 12) cDNA fragment was amplified from NGLY1 lenti ORF clone (Origene, cat. no. RC203447L1, GenBank NM_018297) using PCR primers SA-NGLY1-Ex6-12-F and NGLY1-Ex6-12-R (Supplementary Table 1). The human NGLY1 (exons 6 to 12) cDNA fragment was then cloned into pDONR221 (ThermoFisher) with the following layout (Figure 1A): 5’ homology arm (5’HA) - splice acceptor (SA) - cDNA fragment of exon 6 through exon 12 of human NGLY1 – human growth hormone (hGH) poly(A) signal - floxed mRuby - T2A - puromycin resistance cassette driven by the EF1α promoter - 3’ homology arm (3’HA). The donor vector was designed to integrate into intron 5 of the NGLY1 genomic locus. The 5’ and 3’ HAs were amplified using normal control hiPSC ND2.0 (NIH) genomic DNA as a template. Sequences of corresponding PCR primers (NGLY1 5’HA-F, NGLY1 5’HA-R, NGLY1 3’HA-F, and NGLY1 3’HA-R) for amplifying HAs are listed in Supplementary Table 1. The 5’ and 3’ homology arms (HA) were 488 bp and 600 bp in length, respectively.

As shown in the plasmid layout (**Figure 1A**), the portion of the human NGLY1 cDNA fragment that was comprised of exons 6-12 was preceded by a synthetic splice acceptor (SA) sequence and followed by an hGH polyA sequence. Donor sequences of human NGLY1 were mammalian codon optimized, which have multiple silent mutations that reduce the possibility of premature cross-over events during homology directed recombination (HDR). T2A is the self-cleaving peptide that allows for simultaneous, separate protein expression of puromycin resistance fragment. All fragments were assembled by Gibson assembly (New England Biolabs) as described previously (Gibson et al., 2009; Li et al., 2017). The final construct was selected with kanamycin.

### Construction of a single guide RNA vector for the CRISPR/Cas9 system to target the NGLY1 locus

The pX330-U6-Chimeric_BB-CBh-hSpCas9 vector (Cong et al., 2013), which allows for human single guide RNA (sgRNA) expression, together with mammalian codon-optimized *Streptococcus pyogenes* Cas9 expression, was purchased from Addgene plasmid # 42230; http://n2t.net/addgene:42230; RRID: Addgene_42230). The sequence for making sgRNA for mediating NGLY1 targeting was located within intron 5 of NGLY1: 5’ TAGCTCTTCTCTATATG**GGG** 3’ (PAM sequence is in bold). To construct the sgRNA expression vector, a pair of oligos (NGLY1-f3-top 5’ CACCTAGCTCTTCTCTATATGGGG 3’ and NGLY1-f3-bottom, 5’ AAACCCCCATATAGAGAAGAGCTA3’) was designed using Benchling, a cloud-based biology software (https://www.benchling.com). The oligos were synthesized and subcloned into pX330 sgRNA-Cas9 2-in1-expression vector following a previously published protocol (Ran et al., 2013).

### T7 Endonuclease I (T7EI) assay

T7EI assay was performed as previously described (Guschin et al., 2010; Li et al., 2015; Xue et al., 2016). Briefly, 1.5×10^5^ 293FT cells (one well of a 12-well plate) were transfected with 1 μg NGLY1-sgRNA expression vectors and 1 μg Cas9 expression vector, or GFP (control) vector, using Lipofectamine 3000 (Life Technologies). Cells were harvested 72 h post-transfection, digested with 100 μL genomic DNA quick extraction solution (Fisher), and incubated at 68°C for 15 min, and then 95°C for 8 min. The concentration of the genomic DNA was adjusted to 200 ng/μL before it was PCR amplified using Herculase II (Agilent). Approximately 400 ng purified PCR product (PCR purification kit, Qiagen) was denatured by incubating at 95°C for 5 min, and the temperature was slowly lowered to < 30°C to allow the formation of heteroduplex. T7EI assay reaction was set up as follows: 400 ng hybridized DNA, 2 μL Digestion Buffer 2, and 1 μL (10 units/μL) T7EI enzyme. Reaction was incubated at 37°C for 20 min. Digested product was separated by 2.5% agarose gel. Gel bands were quantified using Image J, and the cleavage efficiency (percentage of insertion and deletion, abbreviated as indel) was calculated following a previously reported protocol (Li *et al*., 2015; Ran *et al*., 2013).

### Generation of gene-corrected isogenic hiPSCs from NGLY1-deficient patient hiPSCs

The NGLY1-deficient hiPSC line was derived from fibroblasts from NGLY1-deficient patients (GM25990) obtained from the Coriell Institute for Medical Research, using the CytoTune Sendai Reprogramming Kit (Thermo Fisher Scientific) in our previous study (Zolekar *et al*., 2018). hiPSCs were maintained under feeder-free conditions on hESC-qualified Matrigel (Corning) in mTeSR Plus medium (Stemcell Technologies) at 37°C and 5% CO_2_. hiPSCs were passaged using ReLeSR (STEM CELL Technologies) once per week. (Li *et al*., 2017; Zheng et al., 2022). Alternatively, cells were passaged every 4∼5 days with Accutase (Innovative Cell Technologies) onto Matrigel (BD Biosciences)-coated culture plates at a ratio of 1:4∼1:8 supplemented with ROCK inhibitor Y-27632 (10 mM, R&D Systems). To monitor and ensure the genetic stability of the cells, routine karyotyping examination was done every 10 to 15 passages. To generate NGLY1 gene-corrected hiPSC lines, approximately 1 × 10^5^ patient hiPSCs were plated in a well of a 12-well plate in mTeSR plus medium, one day prior to transfection. Cells from each well were co-transfected with NGLY1 (0.5 μg) and sgRNA-Cas9 (0.5 μg) vectors using Lipofectamine Stem (Life Technologies) following the manufacturer’s instructions. ROCK inhibitor (10 mM) was added for the first 24 h post-transfection to enhance single-cell survival. Cells were fed with fresh mTeSR plus medium every day until day 3 post-transfection, when puromycin (0.75-1 μg/mL) selection was started. Resistant single clones were isolated and manually picked after two to four days of selection and subsequently expanded individually. Genomic DNA extracted from single clones were examined by PCR for identification of correctly targeted hiPSC clones. Untargeted cells produced a 551-bp PCR product when amplified with the primer pair NGLY1-int5-SurF and NGLY1-int5-SurR. Conversely, successfully targeted (gene-corrected) clones yielded a 949-bp PCR product when amplified with the primer pair, NGLY1-testF1 and NGLY1-qR3 (Primers are listed in Supplementary Table 1). The PCR products with the correct size were verified by Sanger sequencing.

### Differentiation and culture of human iPSC-derived PMPs and microglia

Primitive macrophage progenitors (PMPs) were derived from NGLY1-deficient and CRISPR-mediated, gene corrected hiPSCs using an established yolk sac embryoid body (YS-EB) differentiation protocol (Jiang *et al*., 2020; Jin *et al*., 2026b; Xu *et al*., 2021). To generate YS-EBs, hiPSCs were cultured in mTeSR Plus medium supplemented with BMP4 (50 ng/mL; PeproTech), VEGF (50 ng/mL; PeproTech), and SCF (20 ng/mL; PeproTech) for 6 days. YS-EBs were then transferred to 10-cm dishes containing X-VIVO 15 medium (Lonza) supplemented with IL-3 (25 ng/mL; PeproTech) and M-CSF (100 ng/mL; PeproTech) to induce myeloid differentiation. Floating PMPs emerged in the culture supernatant approximately 4 weeks after plating were collected at matched time points for subsequent analyses. For the microglial differentiation protocol (Jin et al., 2026a; Li et al., 2024), PMPs were cultured for 7 days in Dulbecco’s modified Eagle’s medium (DMEM)/F12 (HyClone) supplemented with N2 (Gibco), IL-34 (100 ng/mL, PeproTech), and GM-CSF (10 ng/mL, PeproTech). NGLY1-deficient and gene corrected cells were differentiated and analyzed in parallel across matched batches.

### Immunocytochemistry

Cultured hiPSCs, PMPs, and hiPSC-derived microglia were fixed with freshly prepared 4% paraformaldehyde in PBS for 15 min at room temperature, washed with PBS, permeabilized with 0.1% Triton X-100 in PBS for 10 min, and blocked with 5% donkey serum–containing blocking buffer for 1 h at room temperature. Cells were then incubated overnight at 4°C with primary antibodies diluted in blocking buffer. All antibodies used are listed in Supplementary Table 2. After incubation with primary antibody, cells were washed with PBS and incubated for 1 h at room temperature with species-appropriate secondary antibodies (Jackson Immuno Research). Nuclei were counterstained with DAPI (Sigma-Aldrich). Samples were mounted in Vectashield and images were captured using a Zeiss AxioVision microscope with z-stack split view function. For comparisons between isogenic pairs or before and after treatments, images were acquired using identical microscope settings within each experiment. Marker-positive cells were quantified relative to total DAPI-positive nuclei unless otherwise stated. RUNX1 nuclear localization was quantified by measuring background-subtracted RUNX1 fluorescence intensity within DAPI-defined nuclei. Microglial morphology was quantified from IBA1-positive cells using ImageJ (Young and Morrison, 2018). Identical thresholding and analysis settings were applied across groups.

### EdU incorporation assay

EdU incorporation was used to assess DNA synthesis in PMPs. Cells were incubated with EdU at a final concentration of 10 μM in culture medium for 2 h at 37°C. After labeling, cells were fixed with freshly prepared 4% paraformaldehyde in PBS for 15 min, washed with PBS, and permeabilized with 0.1% Triton X-100 in PBS for 10 min. EdU detection was performed using a freshly prepared click-reaction solution containing 100 mM Tris-HCl (pH 8.5), 1 mM CuSOΏ, 10 μM Alexa Fluor 647 azide (Thermo Fisher Scientific), and 100 mM ascorbic acid in PBS. Ascorbic acid was added last to initiate the reaction. Cells were incubated with the staining solution for 10-20 min at room temperature in the dark, followed by three washes with PBS. Nuclei were counterstained with DAPI. Images were acquired using identical exposure settings across groups, and EdU-positive cells were quantified as the percentage of EdU-positive nuclei among total DAPI-positive nuclei.

### Western blot

Western blot was performed as described previously (Zhang et al., 2022). Cells were lysed in RIPA buffer supplemented with protease and phosphatase inhibitors (Sigma-Aldrich). Lysates were incubated on ice for 30 min and clarified by centrifugation at 16,000 g for 10min at 4 °C. Protein concentration was determined using *the* Pierce BCA Protein Assay Kit (Thermo Fisher Scientific). Equal amounts of protein, 15–20 μg, were separated on 10% SDS-PAGE gels and transferred to nitrocellulose membranes using a wet transfer system. Membranes were blocked in 5% non-fat milk in TBST for 1 h at room temperature and incubated with primary antibodies overnight at 4 °C. After washing, membranes were incubated with HRP-conjugated secondary antibodies for 1 h at room temperature. Signals were detected using ECL reagents (Pierce ECL Plus Western Blot Substrate, Thermo Scientific) for immunodetection. Band intensities were quantified using ImageJ. O-GlcNAc signals were quantified by total lane intensity after background subtraction and normalized to total protein or loading control. NGLY1, RUNX1, CDKN1C, OGA, and OGT signals were normalized to the loading control.

### Quantitative PCR

Total RNA was extracted from hiPSCs, hiPSC derived-PMPs, or microglia using Quick-RNA miniprep kits (Zymo Research) according to the manufacturer’s instructions, including on-column DNase treatment. RNA concentration and purity were assessed using Nanodrop. Samples with A260/280 ratios between 1.8 and 2.1 were used. cDNA was synthesized from total RNA using the iScript cDNA Synthesis Kit (Bio-Rad). qPCR was performed using the iQ SYBR Green Supermix kit (Bio-Rad). Reactions were run in triplicate with 20 μL per reaction. Cycling conditions were 95 °C, 10 min; 40 cycles of 95 °C, 10 s, and 60 °C, 30 s. The relative fold change in gene expression was calculated using the ΔΔCt method and normalized to GAPDH. Primer sequences are listed in **Supplementary Table S3**.

### GNA measurement in hiPSC derived PMPs

NGLY1-deficient and gene-corrected isogenic PMPs were collected (1 × 10^6^ cells per sample) and cell pellets were processed for LC-MS analysis of aspartylglucosamine (GNA), with a minimum of three independent samples examined for each genotype. Cell pellets were disrupted in 500 μL of extraction buffer composed of acetonitrile:methanol:water (2:2:1), sonicated, vigorously vortexed, and stored for 1 hour at -20°C. Extracts were centrifuged at 21,000 x *g* for 15 minutes and the cleared metabolite-containing extract was transferred to a new tube. A pooled quality-control (QC) sample was generated by combining equal-volume aliquots of metabolite extract from each experimental sample. A 400 μL aliquot of metabolite extract or pooled QC was dried using an Eppendorf Vacufuge Plus without heating. Dried extracts were resuspended in 40 μL of acetonitrile:water (1:1) by vortexing and incubating overnight at -20°C. The following day, samples were centrifuged at 21,000 x *g* for 15 minutes, and the resulting supernatants were transferred to autosampler vials for LC-MS analysis.

For each prepared sample, 2.5 μL was injected onto a SeQuant ZIC-pHILIC column (2.1 X 150 mm, 5 μm particle size, Millipore Sigma cat. no. 150460) fitted with a ZIC-pHILIC guard column (20 x 2.1 mm, Millipore Sigma cat. no. 150437) and coupled to a Thermo Vanquish Flex UHPLC system. The column and autosampler compartments were maintained at 25°C and 4°C, respectively. Mobile phase A consisted of 20 mM ammonium carbonate, 0.1% ammonium hydroxide (v/v), and 10% acetonitrile, and mobile phase B consisted of 90% acetonitrile. Chromatographic separation was performed at a flow rate of 0.150 ml/ min using the following gradient: 0 – 21 min-linear decrease from 87.5 to 12.5% Buffer B; 21 – 21.5 min-linear increase from 12.5 to 87.5% Buffer B; and 21.5–28 min-hold at 87.5% Buffer B.

Data was acquired using a Thermo Exploris 240 mass spectrometer equipped with a heated electrospray ionization (HESI) source. The spray voltage was set to -2.5 kV, the ion-transfer tube temperature to 320°C, and the HESI vaporizer temperature to 350°C. Sheath, auxiliary, and sweep gas flow rates were set to 37, 7, and 0 arbitrary units, respectively. The mass spectrometer was operated in negative-ion full-scan mode over an m/z range of 70 – 1000 at a resolution of 120,000. The AGC target was 2× 10^6^, the maximum injection time was 120 ms, and the RF lens was set to 70%. The QC sample was analyzed at the beginning and at the end of the LC-MS run, as well as intermittently throughout. In addition, the pooled QC was analyzed by negative-ion data-dependent MS/MS (Top10) with stepped HCD collision energies of 10 and 40, an MS/MS resolution of 30,000, an AGC target of 1 × 10^6^, automatic maximum injection time, and a precursor isolation window of 1 m/z to confirm GNA identity. GNA was identified by comparison with an authentic GNA standard (MedChemExpress, cat no. HY-N9446) analyzed under the same chromatographic and MS/MS conditions. GNA was detected as the [M−H]− ion at m/z 334.1256 with a retention time of approximately 10.1 min. Identity was further confirmed by its MS/MS fragmentation spectrum, which contained product ions at m/z 316.114, 113.035, 214.083, 196.073, and 96.009, listed in decreasing order of intensity.

Raw integrated peak areas were obtained for all experimental and QC samples using Thermo TraceFinder 5.3 set to a mass tolerance of ± 2.0 mmu. Analytical signal drift was corrected using QC-MXP (QCMXP, v3.0) with default settings, using repeated pooled-QC measurements to model and correct changes in signal intensity over the analytical sequence. To normalize for sample-to-sample variation, the QC-corrected peak area from 180 additional metabolite features was used to generate a sample-specific relative scaling factor. For each feature, the corrected peak area was divided by its median value across all experimental samples. The resulting median-scaled values were summed within each sample, and each sum was divided by the median sum across the complete dataset to generate a sample-specific relative scaling factor. QC-corrected GNA peak areas were then divided by the corresponding scaling factor to obtain normalized relative GNA abundance. Fold changes were calculated using the appropriate control group as the reference.

### Pharmacological modulation of O-GlcNAc cycling

To modulate O-GlcNAc cycling, PMPs were treated with the OGT inhibitor OSMI-1 and the OGA inhibitor Thiamet-G (MedChemExpress, MCE). Cells were treated with OSMI-1 concentration 0, 5, 10, or 20 μM or Thiamet-G concentration 0, 2, 5, or 10 μM for 24 h. Vehicle-treated cells received DMSO at the same final concentration. Dose-response experiments were performed to identify concentrations that altered total O-GlcNAcylation without overt toxicity. Based on these responses, we selected 5 μM Thiamet-G and 10 μM OSMI-1 for subsequent treatment of gene corrected and NGLY1-deficient PMPs. Cells were collected for Western blot, immunofluorescence staining, and qPCR.

### LPS stimulation and inflammatory response assay

To assess inflammatory responsiveness, iPSC-derived microglia were treated with LPS (Sigma-Aldrich) at 0, 0.1, or 1 μM for 24 h. After stimulation, cells underwent pHrodo assay for phagocytosis, immunocytochemistry staining for CD68 or IBA1, or were collected for RNA extraction and subsequent qPCR analysis of expression of *TNFα*, *IL1B*, and *IL6*.

### Phagocytosis assay

Phagocytic capacity was assessed using pHrodo bioparticles (Thermo Fisher Scientific). NGLY1-deficient and gene corrected iPSC-derived microglia were seeded in 96-well plates at 1 × 10^4^ cells per well. Where indicated, cells were treated with LPS at 0, 0.1, or 1 μM before the assay. Nuclei were stained with DAPI. pHrodo particles were vortexed, ultrasonicated for 5 min, diluted in live-cell imaging solution (Thermo Fisher Scientific), and added to live microglia. Cells were incubated for 2 h at 37°C, washed to remove non-engulfed particles, and imaged using a Keyence BZ-X800 microscope. Phagocytosis was quantified as the percentage of pHrodo-positive microglia using identical analysis parameters across groups.

### RUNX1 knockdown using shRNA lentiviral vectors

RUNX1 knockdown was performed in undifferentiated NGLY1-deficient hiPSCs using lentiviral expression vectors that carry independent RUNX1 targeting shRNA sequences: RUNX1 shRNA (H749) or RUNX1 shRNA (H750). Non-targeting (NT) shRNA (H748) was used in the control group. hiPSCs were transduced overnight in mTeSR Plus medium, and replaced with fresh medium the following day. After 72 h, cells were selected with puromycin (1 μg/mL) for two days. Stable hiPSCs were then used for PMP differentiation. Knockdown efficiency was confirmed by examining RUNX1 expression at both mRNA and protein levels (**Figure 5**).

### RNA library construction, quality control, sequencing, and bioinformatic analysis

Total RNA was extracted from hiPSC-derived PMPs using the Quick-RNA miniprep kits (Zymo Research) according to the manufacturer’s instructions. RNA concentration, purity, and integrity were assessed using NanoDrop and an Agilent Bioanalyzer. Samples with RIN ≥ 6.8 were used for library preparation and sequencing (Novogene). Reads were aligned to the human reference genome hg38 using HISAT2 v2.1.0, and gene-level counts were generated using featureCounts from the Rsubread package. Differential expression analysis was performed using DESeq2, with p-values adjusted using the Benjamini-Hochberg method. Gene Ontology and GSEA analyses were performed to assess pathway-level changes.

### Statistical analysis

Statistical analyses were performed using GraphPad Prism version 10.0 (GraphPad Software, Boston, MA) and R, where indicated. Data are presented as mean ± SEM unless stated otherwise. Biological replicates were defined as independent differentiations initiated from separate hiPSC cultures or passages on different days. For imaging-based assays, multiple cells or fields from the same differentiation were averaged before statistical testing unless otherwise specified. For comparisons between two independent groups, two-tailed unpaired t tests with Welch’s correction were used. For matched comparisons between NGLY1-deficient and gene corrected cells generated within the same differentiation batch, paired tests were used where appropriate. For experiments involving multiple groups or treatment conditions, two-way ANOVA followed by Tukey’s multiple-comparison tests were used, as indicated in the figure legends. A p-value < 0.05 was considered significant. Significance is indicated as \**p* < 0.05, \*\**p* < 0.01, \*\*\**p* < 0.001, NS, not significant. Unless otherwise noted, experiments were independently performed at least three times with similar results.

## Acknowledgments

This work was supported by FIU start-up funds (YL).

## Author Contributions

Conceptualization, HZ and YL; Methodology, HZ, HX, AJR, YCW, and YL; Investigation, HZ, HX, MJ, TW, AJR, PJ, YCW, and YL; Writing – Original Draft, HZ, AJR, and YL; Writing – Review & Editing, HZ, HX, MJ, TW, AJR, PJ, YCW, and YL; Funding Acquisition, YL; Resources, AJR, PJ, YCW, and YL; Supervision, AJR, PJ, YCW, and YL.

## References

Abbott, J., Tambe, M., Pavlinov, I., Farkhondeh, A., Nguyen, H.N., Xu, M., Pradhan, M., York, T., Might, M., Baumgärtel, K., et al. (2023). Generation and characterization of NGLY1 patient-derived midbrain organoids. Front Cell Dev Biol 11, 1039182. 10.3389/fcell.2023.1039182.

Abud, E.M., Ramirez, R.N., Martinez, E.S., Healy, L.M., Nguyen, C.H.H., Newman, S.A., Yeromin, A.V., Scarfone, V.M., Marsh, S.E., Fimbres, C., et al. (2017). iPSC-Derived Human Microglia-like Cells to Study Neurological Diseases. Neuron 94, 278–293.e279. 10.1016/j.neuron.2017.03.042.

Asahina, M., Fujinawa, R., Fujihira, H., Masahara-Negishi, Y., Andou, T., Tozawa, R., and Suzuki, T. (2021a). JF1/B6F1 Ngly1(-/-) mouse as an isogenic animal model of NGLY1 deficiency. Proc Jpn Acad Ser B Phys Biol Sci 97, 89–102. 10.2183/pjab.97.005.

Asahina, M., Fujinawa, R., Hirayama, H., Tozawa, R., Kajii, Y., and Suzuki, T. (2021b). Reversibility of motor dysfunction in the rat model of NGLY1 deficiency. Mol Brain 14, 91. 10.1186/s13041-021-00806-6.

Asahina, M., Fujinawa, R., Nakamura, S., Yokoyama, K., Tozawa, R., and Suzuki, T. (2020). Ngly1 -/- rats develop neurodegenerative phenotypes and pathological abnormalities in their peripheral and central nervous systems. Hum Mol Genet 29, 1635–1647. 10.1093/hmg/ddaa059.

Borges, K.S., Arboleda, V.A., and Vilain, E. (2015). Mutations in the PCNA-binding site of CDKN1C inhibit cell proliferation by impairing the entry into S phase. Cell Div 10, 2. 10.1186/s13008-015-0008-8.

Butovsky, O., Jedrychowski, M.P., Moore, C.S., Cialic, R., Lanser, A.J., Gabriely, G., Koeglsperger, T., Dake, B., Wu, P.M., Doykan, C.E., et al. (2014). Identification of a unique TGF-β-dependent molecular and functional signature in microglia. Nat Neurosci 17, 131–143. 10.1038/nn.3599.

Caglayan, A.O., Comu, S., Baranoski, J.F., Parman, Y., Kaymakçalan, H., Akgumus, G.T., Caglar, C., Dolen, D., Erson-Omay, E.Z., Harmanci, A.S., et al. (2015). NGLY1 mutation causes neuromotor impairment, intellectual disability, and neuropathy. European Journal of Medical Genetics 58, 39–43. 10.1016/j.ejmg.2014.08.008.

Chou, T.Y., Dang, C.V., and Hart, G.W. (1995). Glycosylation of the c-Myc transactivation domain. Proc Natl Acad Sci U S A 92, 4417–4421. 10.1073/pnas.92.10.4417.

Cong, L., Ran, F.A., Cox, D., Lin, S., Barretto, R., Habib, N., Hsu, P.D., Wu, X., Jiang, W., Marraffini, L.A., and Zhang, F. (2013). Multiplex genome engineering using CRISPR/Cas systems. Science 339, 819–823. 10.1126/science.1231143.

Du, A., Yang, K., Zhou, X., Ren, L., Liu, N., Zhou, C., Liang, J., Yan, N., Gao, G., and Wang, D. (2024). Systemic gene therapy corrects the neurological phenotype in a mouse model of NGLY1 deficiency. JCI Insight 9. 10.1172/jci.insight.183189.

Enns, G.M., Shashi, V., Bainbridge, M., Gambello, M.J., Zahir, F.R., Bast, T., Crimian, R., Schoch, K., Platt, J., Cox, R., et al. (2014). Mutations in NGLY1 cause an inherited disorder of the endoplasmic reticulum-associated degradation pathway. Genet Med 16, 751–758. 10.1038/gim.2014.22.

Friedman, B.A., Srinivasan, K., Ayalon, G., Meilandt, W.J., Lin, H., Huntley, M.A., Cao, Y., Lee, S.H., Haddick, P.C.G., Ngu, H., et al. (2018). Diverse Brain Myeloid Expression Profiles Reveal Distinct Microglial Activation States and Aspects of Alzheimer’s Disease Not Evident in Mouse Models. Cell Rep 22, 832–847. 10.1016/j.celrep.2017.12.066.

Galatro, T.F., Holtman, I.R., Lerario, A.M., Vainchtein, I.D., Brouwer, N., Sola, P.R., Veras, M.M., Pereira, T.F., Leite, R.E.P., Möller, T., et al. (2017). Transcriptomic analysis of purified human cortical microglia reveals age-associated changes. Nat Neurosci 20, 1162–1171. 10.1038/nn.4597.

Geirsdottir, L., David, E., Keren-Shaul, H., Weiner, A., Bohlen, S.C., Neuber, J., Balic, A., Giladi, A., Sheban, F., Dutertre, C.A., et al. (2019). Cross-Species Single-Cell Analysis Reveals Divergence of the Primate Microglia Program. Cell 179, 1609–1622 e1616. 10.1016/j.cell.2019.11.010.

Gibson, D.G., Young, L., Chuang, R.Y., Venter, J.C., Hutchison, C.A., 3rd, and Smith, H.O. (2009). Enzymatic assembly of DNA molecules up to several hundred kilobases. Nat Methods 6, 343–345. 10.1038/nmeth.1318.

Ginhoux, F., Greter, M., Leboeuf, M., Nandi, S., See, P., Gokhan, S., Mehler, M.F., Conway, S.J., Ng, L.G., Stanley, E.R., et al. (2010). Fate mapping analysis reveals that adult microglia derive from primitive macrophages. Science 330, 841–845. 10.1126/science.1194637.

Gosselin, D., Skola, D., Coufal, N.G., Holtman, I.R., Schlachetzki, J.C.M., Sajti, E., Jaeger, B.N., O’Connor, C., Fitzpatrick, C., Pasillas, M.P., et al. (2017). An environment-dependent transcriptional network specifies human microglia identity. Science 356. 10.1126/science.aal3222.

Guschin, D.Y., Waite, A.J., Katibah, G.E., Miller, J.C., Holmes, M.C., and Rebar, E.J. (2010). A rapid and general assay for monitoring endogenous gene modification. Methods Mol Biol 649, 247–256. 10.1007/978-1-60761-753-2_15.

Guttikonda, S.R., Sikkema, L., Tchieu, J., Saurat, N., Walsh, R.M., Harschnitz, O., Ciceri, G., Sneeboer, M., Mazutis, L., Setty, M., et al. (2021). Fully defined human pluripotent stem cell-derived microglia and tri-culture system model C3 production in Alzheimer’s disease. Nat Neurosci 24, 343–354. 10.1038/s41593-020-00796-z.

Haenseler, W., Sansom, S.N., Buchrieser, J., Newey, S.E., Moore, C.S., Nicholls, F.J., Chintawar, S., Schnell, C., Antel, J.P., Allen, N.D., et al. (2017). A Highly Efficient Human Pluripotent Stem Cell Microglia Model Displays a Neuronal-Co-culture-Specific Expression Profile and Inflammatory Response. Stem Cell Reports 8, 1727–1742. 10.1016/j.stemcr.2017.05.017.

Haijes, H.A., de Sain-van der Velden, M.G.M., Prinsen, H., Willems, A.P., van der Ham, M., Gerrits, J., Couse, M.H., Friedman, J.M., van Karnebeek, C.D.M., Selby, K.A., et al. (2019). Aspartylglycosamine is a biomarker for NGLY1-CDDG, a congenital disorder of deglycosylation. Mol Genet Metab 127, 368–372. 10.1016/j.ymgme.2019.07.001.

Han, I., and Kudlow, J.E. (1997). Reduced O glycosylation of Sp1 is associated with increased proteasome susceptibility. Mol Cell Biol 17, 2550–2558. 10.1128/MCB.17.5.2550.

Hart, G.W., Slawson, C., Ramirez-Correa, G., and Lagerlof, O. (2011). Cross talk between O-GlcNAcylation and phosphorylation: roles in signaling, transcription, and chronic disease. Annu Rev Biochem 80, 825–858. 10.1146/annurev-biochem-060608-102511.

Housley, M.P., Rodgers, J.T., Udeshi, N.D., Kelly, T.J., Shabanowitz, J., Hunt, D.F., Puigserver, P., and Hart, G.W. (2008). O-GlcNAc regulates FoxO activation in response to glucose. J Biol Chem 283, 16283–16292. 10.1074/jbc.M802240200.

Huang, C., Harada, Y., Hosomi, A., Masahara-Negishi, Y., Seino, J., Fujihira, H., Funakoshi, Y., Suzuki, T., Dohmae, N., and Suzuki, T. (2015). Endo-β-N-acetylglucosaminidase forms N-GlcNAc protein aggregates during ER-associated degradation in Ngly1-defective cells. Proc Natl Acad Sci U S A 112, 1398–1403. 10.1073/pnas.1414593112.

Hubschmann, V., Korkut-Demirbas, M., and Siegert, S. (2022). Assessing human iPSC-derived microglia identity and function by immunostaining, phagocytosis, calcium activity, and inflammation assay. STAR Protoc 3, 101866. 10.1016/j.xpro.2022.101866.

Jiang, P., Turkalj, L., and Xu, R. (2020). High-Fidelity Modeling of Human Microglia with Pluripotent Stem Cells. Cell Stem Cell 26, 629–631. 10.1016/j.stem.2020.04.004.

Jin, M., Ma, Z., Dang, R., Zhang, H., Kim, R., Xue, H., Pascual, J., Yu, H., Papetti, A.V., Liu, Y., et al. (2026a). A myeloid trisomy 21-associated gene variant is protective from Alzheimer’s disease. Nat Neurosci 29, 25–39. 10.1038/s41593-025-02117-8.

Jin, M., Ma, Z., Zhang, H., Dang, R., Papetti, A.V., Stillitano, A.C., Zou, L., Goldman, S.A., and Jiang, P. (2026b). Chimeric brain models to study human glial-neuronal and macroglial-microglial interactions. Cell Rep 45, 116794. 10.1016/j.celrep.2025.116794.

Lam, C., Ferreira, C., Krasnewich, D., Toro, C., Latham, L., Zein, W.M., Lehky, T., Brewer, C., Baker, E.H., Thurm, A., et al. (2017). Prospective phenotyping of NGLY1-CDDG, the first congenital disorder of deglycosylation. Genetics in Medicine 19, 160–168. 10.1038/gim.2016.75.

Lancrin, C., Sroczynska, P., Stephenson, C., Allen, T., Kouskoff, V., and Lacaud, G. (2009). The haemangioblast generates haematopoietic cells through a haemogenic endothelium stage. Nature 457, 892–895. 10.1038/nature07679.

Lehrbach, N.J., Breen, P.C., and Ruvkun, G. (2019). Protein Sequence Editing of SKN-1A/Nrf1 by Peptide:N-Glycanase Controls Proteasome Gene Expression. Cell 177, 737–750.e715. 10.1016/j.cell.2019.03.035.

Li, S., Xue, H., Long, B., Sun, L., Truong, T., and Liu, Y. (2015). Efficient generation of hiPSC neural lineage specific knockin reporters using the CRISPR/Cas9 and Cas9 double nickase system. J Vis Exp, e52539. 10.3791/52539.

Li, S., Zhang, A., Xue, H., Li, D., and Liu, Y. (2017). One-Step piggyBac Transposon-Based CRISPR/Cas9 Activation of Multiple Genes. Mol Ther Nucleic Acids 8, 64–76. 10.1016/j.omtn.2017.06.007.

Li, X., Liu, J., Boreland, A.J., Kapadia, S., Zhang, S., Stillitano, A.C., Abbo, Y., Clark, L., Lai, D., Liu, Y., et al. (2024). Polygenic risk for alcohol use disorder affects cellular responses to ethanol exposure in a human microglial cell model. Sci Adv 10, eado5820. 10.1126/sciadv.ado5820.

Lin, V.J.T., Hu, J., Zolekar, A., Salick, M.R., Mittal, P., Bird, J.T., Hoffmann, P., Kaykas, A., Byrum, S.D., and Wang, Y.C. (2022). Deficiency of N-glycanase 1 perturbs neurogenesis and cerebral development modeled by human organoids. Cell Death Dis 13, 262. 10.1038/s41419-022-04693-0.

Manole, A., Wong, T., Rhee, A., Novak, S., Chin, S.M., Tsimring, K., Paucar, A., Williams, A., Newmeyer, T.F., Schafer, S.T., et al. (2023). NGLY1 mutations cause protein aggregation in human neurons. Cell Rep 42, 113466. 10.1016/j.celrep.2023.113466.

Matsuoka, S., Edwards, M.C., Bai, C., Parker, S., Zhang, P., Baldini, A., Harper, J.W., and Elledge, S.J. (1995). p57KIP2, a structurally distinct member of the p21CIP1 Cdk inhibitor family, is a candidate tumor suppressor gene. Genes Dev 9, 650–662. 10.1101/gad.9.6.650.

Mesika, A., Nadav, G., Shochat, C., Kalfon, L., Jackson, K., Khalaileh, A., Karasik, D., and Falik-Zaccai, T.C. (2022). NGLY1 Deficiency Zebrafish Model Manifests Abnormalities of the Nervous and Musculoskeletal Systems. Front Cell Dev Biol 10, 902969. 10.3389/fcell.2022.902969.

Mueller, W.F., Jakob, P., Sun, H., Clauder-Münster, S., Ghidelli-Disse, S., Ordonez, D., Boesche, M., Bantscheff, M., Collier, P., Haase, B., et al. (2020). Loss of N-Glycanase 1 Alters Transcriptional and Translational Regulation in K562 Cell Lines. G3: Genes Genomes Genetics (Bethesda) 10, 1585–1597. 10.1534/g3.119.401031.

Mueller, W.F., Zhu, L., Tan, B., Dwight, S., Beahm, B., Wilsey, M., Wechsler, T., Mak, J., Cowan, T., Pritchett, J., et al. (2022). GlcNAc-Asn is a biomarker for NGLY1 deficiency. J Biochem 171, 177–186. 10.1093/jb/mvab111.

Muffat, J., Li, Y., Yuan, B., Mitalipova, M., Omer, A., Corcoran, S., Bakiasi, G., Tsai, L.H., Aubourg, P., Ransohoff, R.M., and Jaenisch, R. (2016). Efficient derivation of microglia-like cells from human pluripotent stem cells. Nat Med 22, 1358–1367. 10.1038/nm.4189.

Na, H.J., Abramowitz, L.K., and Hanover, J.A. (2022). Cytosolic O-GlcNAcylation and PNG1 maintain Drosophila gut homeostasis by regulating proliferation and apoptosis. PLoS Genet 18, e1010128. 10.1371/journal.pgen.1010128.

Need, A.C., Shashi, V., Hitomi, Y., Schoch, K., Shianna, K.V., McDonald, M.T., Meisler, M.H., and Goldstein, D.B. (2012). Clinical application of exome sequencing in undiagnosed genetic conditions. J Med Genet 49, 353–361. 10.1136/jmedgenet-2012-100819.

Padmakumar, M., Biesmans, S., Valadas, J.S., Detrez, J.R., Gillet, G., Bresler, P., Clénet, M.-L., and Kadiu, I. (2023). An improved method for large scale generation and high-throughput functional characterization of human iPSC-derived microglia. Volume 3 -2023. 10.3389/fddsv.2023.1289314.

Pandey, A., Adams, J.M., Han, S.Y., and Jafar-Nejad, H. (2022). NGLY1 Deficiency, a Congenital Disorder of Deglycosylation: From Disease Gene Function to Pathophysiology. Cells 11. 10.3390/cells11071155.

Paolicelli, R.C., Bolasco, G., Pagani, F., Maggi, L., Scianni, M., Panzanelli, P., Giustetto, M., Ferreira, T.A., Guiducci, E., Dumas, L., et al. (2011). Synaptic pruning by microglia is necessary for normal brain development. Science 333, 1456–1458. 10.1126/science.1202529.

Parker, M.P., Peterson, K.R., and Slawson, C. (2021). O-GlcNAcylation and O-GlcNAc Cycling Regulate Gene Transcription: Emerging Roles in Cancer. Cancers (Basel) 13. 10.3390/cancers13071666.

Prinz, M., Jung, S., and Priller, J. (2019). Microglia Biology: One Century of Evolving Concepts. Cell 179, 292–311. 10.1016/j.cell.2019.08.053.

Ran, F.A., Hsu, P.D., Wright, J., Agarwala, V., Scott, D.A., and Zhang, F. (2013). Genome engineering using the CRISPR-Cas9 system. Nat Protoc 8, 2281–2308. 10.1038/nprot.2013.143.

Salic, A., and Mitchison, T.J. (2008). A chemical method for fast and sensitive detection of DNA synthesis in vivo. Proc Natl Acad Sci U S A 105, 2415–2420. 10.1073/pnas.0712168105.

Schafer, D.P., Lehrman, E.K., Kautzman, A.G., Koyama, R., Mardinly, A.R., Yamasaki, R., Ransohoff, R.M., Greenberg, M.E., Barres, B.A., and Stevens, B. (2012). Microglia sculpt postnatal neural circuits in an activity and complement-dependent manner. Neuron 74, 691–705. 10.1016/j.neuron.2012.03.026.

Sekine, H., and Motohashi, H. (2021). Roles of CNC Transcription Factors NRF1 and NRF2 in Cancer. Cancers (Basel) 13. 10.3390/cancers13030541.

Shyr, Z.A., Amniouel, S., Owusu-Ansah, K., Tambe, M., Abbott, J., Might, M., and Zheng, W. (2025). Increased oxidative stress and autophagy in NGLY1 patient iPSC-derived neural stem cells. Exp Cell Res 448, 114540. 10.1016/j.yexcr.2025.114540.

Sierra, A., Encinas, J.M., Deudero, J.J., Chancey, J.H., Enikolopov, G., Overstreet-Wadiche, L.S., Tsirka, S.E., and Maletic-Savatic, M. (2010). Microglia shape adult hippocampal neurogenesis through apoptosis-coupled phagocytosis. Cell Stem Cell 7, 483–495. 10.1016/j.stem.2010.08.014.

Suzuki, T. (2015). The cytoplasmic peptide:N-glycanase (Ngly1)-basic science encounters a human genetic disorder. J Biochem 157, 23–34. 10.1093/jb/mvu068.

Suzuki, T., Huang, C., and Fujihira, H. (2016). The cytoplasmic peptide:N-glycanase (NGLY1) - Structure, expression and cellular functions. Gene 577, 1–7. 10.1016/j.gene.2015.11.021.

Tomlin, F.M., Gerling-Driessen, U.I.M., Liu, Y.C., Flynn, R.A., Vangala, J.R., Lentz, C.S., Clauder-Muenster, S., Jakob, P., Mueller, W.F., Ordoñez-Rueda, D., et al. (2017). Inhibition of NGLY1 Inactivates the Transcription Factor Nrf1 and Potentiates Proteasome Inhibitor Cytotoxicity. ACS Cent Sci 3, 1143–1155. 10.1021/acscentsci.7b00224.

Tong, S., Ventola, P., Frater, C.H., Klotz, J., Phillips, J.M., Muppidi, S., Dwight, S.S., Mueller, W.F., Beahm, B.J., Wilsey, M., and Lee, K.J. (2023). NGLY1 deficiency: a prospective natural history study. Hum Mol Genet 32, 2787–2796. 10.1093/hmg/ddad106.

Xing, X., Wang, H., Niu, T., Jiang, Y., Shi, X., and Liu, K. (2021). RUNX1 can mediate the glucose and O-GlcNAc-driven proliferation and migration of human retinal microvascular endothelial cells. BMJ Open Diabetes Res Care 9. 10.1136/bmjdrc-2020-001898.

Xu, R., Boreland, A.J., Li, X., Erickson, C., Jin, M., Atkins, C., Pang, Z.P., Daniels, B.P., and Jiang, P. (2021). Developing human pluripotent stem cell-based cerebral organoids with a controllable microglia ratio for modeling brain development and pathology. Stem Cell Reports 16, 1923–1937. 10.1016/j.stemcr.2021.06.011.

Xu, R., Li, X., Boreland, A.J., Posyton, A., Kwan, K., Hart, R.P., and Jiang, P. (2020). Human iPSC-derived mature microglia retain their identity and functionally integrate in the chimeric mouse brain. Nature Communications 11, 1577. 10.1038/s41467-020-15411-9.

Xue, H., Wu, J., Li, S., Rao, M.S., and Liu, Y. (2016). Genetic Modification in Human Pluripotent Stem Cells by Homologous Recombination and CRISPR/Cas9 System. Methods Mol Biol 1307, 173–190. 10.1007/7651_2014_73.

Yang, K., Huang, R., Fujihira, H., Suzuki, T., and Yan, N. (2018). N-glycanase NGLY1 regulates mitochondrial homeostasis and inflammation through NRF1. Journal of Experimental Medicine 215, 2600–2616. 10.1084/jem.20180783.

Yang, K., Torres-Ramirez, G., Dobbs, N., Han, J., Asahina, M., Fujinawa, R., Song, K., Liu, Y., Lin, W., Oviedo, A., et al. (2025). The STING pathway drives noninflammatory neurodegeneration in NGLY1 deficiency. J Exp Med 222. 10.1084/jem.20242296.

Yang, W.H., Kim, J.E., Nam, H.W., Ju, J.W., Kim, H.S., Kim, Y.S., and Cho, J.W. (2006). Modification of p53 with O-linked N-acetylglucosamine regulates p53 activity and stability. Nat Cell Biol 8, 1074–1083. 10.1038/ncb1470.

Yang, W.H., Park, S.Y., Nam, H.W., Kim, D.H., Kang, J.G., Kang, E.S., Kim, Y.S., Lee, H.C., Kim, K.S., and Cho, J.W. (2008). NFkappaB activation is associated with its O-GlcNAcylation state under hyperglycemic conditions. Proc Natl Acad Sci U S A 105, 17345–17350. 10.1073/pnas.0806198105.

Yang, X., and Qian, K. (2017). Protein O-GlcNAcylation: emerging mechanisms and functions. Nat Rev Mol Cell Biol 18, 452–465. 10.1038/nrm.2017.22.

Yang, X., Su, K., Roos, M.D., Chang, Q., Paterson, A.J., and Kudlow, J.E. (2001). O-linkage of N-acetylglucosamine to Sp1 activation domain inhibits its transcriptional capability. Proc Natl Acad Sci U S A 98, 6611–6616. 10.1073/pnas.111099998.

Young, K., and Morrison, H. (2018). Quantifying Microglia Morphology from Photomicrographs of Immunohistochemistry Prepared Tissue Using ImageJ. J Vis Exp. 10.3791/57648.

Zhang, H., Li, H., Lu, M., Wang, S., Ma, X., Wang, F., Liu, J., Li, X., Yang, H., Zhang, F., et al. (2022). Repressor element 1-silencing transcription factor deficiency yields profound hearing loss through K(v)7.4 channel upsurge in auditory neurons and hair cells. Elife 11. 10.7554/eLife.76754.

Zhang, H., Xue, H., Wang, Y.C., and Liu, Y. (2025). NGLY1 as an Emerging Critical Modulator for Neurodevelopment and Pathogenesis in the Brain. Int J Mol Sci 26. 10.3390/ijms26199705.

Zheng, Y., Gallegos, C.M., Xue, H., Li, S., Kim, D.H., Zhou, H., Xia, X., Liu, Y., and Cao, Q. (2022). Transplantation of Human Induced Pluripotent Stem Cell-Derived Neural Progenitor Cells Promotes Forelimb Functional Recovery after Cervical Spinal Cord Injury. Cells 11. 10.3390/cells11172765.

Zhou, S., Zhang, Y., Belmar, J., Hou, C., Zhang, Y., Peng, C., Meng, Y., Li, Z., Mughal, M.J., Gao, Y., et al. (2025). Stabilization of RUNX1 Induced by O-GlcNAcylation Promotes PDGF-BB-Mediated Resistance to CDK4/6 Inhibitors in Breast Cancer. Cancer Res 85, 1708–1724. 10.1158/0008-5472.Can-24-2492.

Zhu, L., Tan, B., Dwight, S.S., Beahm, B., Wilsey, M., Crawford, B.E., Schweighardt, B., Cook, J.W., Wechsler, T., and Mueller, W.F. (2022). AAV9-NGLY1 gene replacement therapy improves phenotypic and biomarker endpoints in a rat model of NGLY1 Deficiency. Mol Ther Methods Clin Dev 27, 259–271. 10.1016/j.omtm.2022.09.015.

Zolekar, A., Lin, V.J.T., Mishra, N.M., Ho, Y.Y., Hayatshahi, H.S., Parab, A., Sampat, R., Liao, X., Hoffmann, P., Liu, J., et al. (2018). Stress and interferon signalling-mediated apoptosis contributes to pleiotropic anticancer responses induced by targeting NGLY1. Br J Cancer 119, 1538–1551. 10.1038/s41416-018-0265-9.

