## supplementary for "An isogenic human stem cell model reveals NGLY1-dependent control of microglial development"

### Supplementary Figure Legends

**Supplementary Figure S1. Verification and basic characterization of NGLY1-deficient and gene-corrected isogenic hiPSCs**.

**(A)** Identification of homozygous gene-corrected hiPSC clones following CRISPR-mediated gene correction of NGLY1-deficient patient hiPSCs. Genomic DNA extracted from individual clones was analyzed by PCR amplification of the targeted NGLY1 genomic locus. Clones correctly targeted at both alleles showed a 949-bp targeted band with the absence of the 551-bp WT band (also see **Figure 1A** for the PCR design). Correctly targeted clones No. 7, 8, and 17 are indicated by asterisks (*). Gene correction at the NGLY1 loci in exons 8 and 11 was further verified by Sanger sequencing of the PCR products (**Figure 1B**).

**(B)** Representative images showing expression of pluripotency markers OCT4, TRA1-81, and NANOG in NGLY1-deficient and gene-corrected hiPSCs. Scale bars, 100 µm.

**(C)** G-band karyotyping analysis of NGLY1-deficient patient hiPSCs and a gene-corrected isogenic hiPSC clone.

**Supplementary Figure S2. Pharmacological modulation of O-GlcNAc cycling in PMPs.**

**(A)** Western blot analysis of NGLY1 protein expression in NGLY1-deficient and gene-corrected PMPs. NGLY1 expression was restored in gene-corrected hiPSC-derived PMPs but was significantly lower in NGLY1-deficient PMPs. Three independent experiments (n = 3) were performed. Data are presented as mean ± SEM.

**(B)** Western blot analysis of CDKN1C ($p57^{kip2}$) protein expression in NGLY1-deficient and gene-corrected PMPs. CDKN1C was significantly increased in NGLY1-deficient PMPs compared with gene-corrected PMPs, consistent with the corresponding mRNA expression levels (see **Figure 2D**). Three independent experiments (n = 3) were performed. Data are presented as mean ± SEM.

**(C)** Dose-response analysis of global O-GlcNAcylation in PMPs derived from a normal hiPSC line (WT control PMPs) following treatment with the OGA inhibitor Thiamet-G or the OGT inhibitor OSMI-1 for 24 h. Thiamet-G was applied at 0, 2, 5, or 10 µM, and OSMI-1 at 0, 5, 10, or 20 µM. Global O-GlcNAcylation levels were assessed by Western blot analysis using the RL2 antibody. GAPDH was used as a loading control. Three independent experiments (n = 3) were performed. Data are presented as mean ± SEM.

**(D)** Western blot analysis of global O-GlcNAcylation in NGLY1-deficient and gene-corrected PMPs after treatment with OSMI-1 or Thiamet-G. NGLY1-deficient PMPs were treated with OSMI-1 at 10 µM for 24 h, and gene-corrected isogenic PMPs were treated with Thiamet-G at 5 µM for 24 h. Global O-GlcNAcylation levels were assessed by Western blot analysis using the RL2 antibody. GAPDH was used as a loading control. Three independent experiments (n = 3) were performed. Data are presented as mean ± SEM.

**(E)** Western blot analysis of RUNX1 protein expression in NGLY1-deficient and gene-corrected PMPs after treatment with OSMI-1 or Thiamet-G. NGLY1-deficient PMPs were treated with OSMI-1 at 10 µM for 24 h, and gene-corrected isogenic PMPs were treated with Thiamet-G at 5 µM for 24 h. GAPDH was used as a loading control. Three independent experiments (n = 3) were performed. Data are presented as mean ± SEM.

**Supplementary Figure S3. Extended exploratory transcriptomic analyses of NGLY1-deficient PMPs**

**(A)** GO cellular component and molecular function enrichment analysis of genes upregulated in NGLY1-deficient PMPs relative to gene-corrected isogenic PMPs. Selected cellular component and molecular function terms are shown. Bars represent −log10 adjusted p-value. CC, cellular component; MF, molecular function.

**(B)** GO cellular component and molecular function enrichment analysis of genes downregulated in NGLY1-deficient PMPs relative to gene-corrected isogenic PMPs. Selected cellular component and molecular function terms are shown. Bars represent −log10 adjusted p-value. CC, cellular component; MF, molecular function.

**(C)** Heatmap of selected immune-, inflammatory-, myeloid-, and progenitor-associated transcripts in NGLY1-deficient and gene-corrected isogenic PMPs. Expression values are displayed as row-wise Z-scores of relative expressions.

**(D)** Gene set enrichment plot for the N-glycan trimming in the endoplasmic reticulum and calnexin/calreticulin cycle gene set.

**(E)** Gene set enrichment plot for the G1/S DNA damage checkpoint gene set.

**Supplementary Figure S4. Validation of RUNX1 shRNA constructs and differentiation capacity of RUNX1 knockdown hiPSCs**

**(A)** Schematic of the lentiviral shRNA expression vector used for stable RUNX1 knockdown. The vector contains the 5′ LTR, EF1α promoter, miR30-based shRNA cassette, WPRE element, puromycin resistance cassette, and 3′ LTR. The sequences and hairpin structures of two independent RUNX1-targeting shRNAs, RUNX1 shRNA749 and RUNX1 shRNA750, are shown.

**(B)** Representative images of OCT4 and TRA1-81 in NGLY1-deficient hiPSCs expressing NT shRNA control, RUNX1 shRNA749, or RUNX1 shRNA750. Scale bars, 50 μm.

**(C)** Representative images of CD43 and CD235 in PMPs differentiated from NGLY1-deficient hiPSCs expressing NT shRNA control, RUNX1 shRNA749, or RUNX1 shRNA750. Scale bars, 20 μm.

**(D)** Representative images of IBA1 in iPSC-derived microglia generated from NGLY1-deficient hiPSCs expressing NT shRNA control, RUNX1 shRNA749, or RUNX1 shRNA750. Scale bars, 50 μm.

**Supplementary Table S1:** PCR primer sequences

| Name | Sequence (5'-3') |
|---|---|
| **attL1-h NGLY1-int5-5HA-F** | CTTTGTACAAAAAAGCAGGCTTAGTTCTAAAGTATAACTTGAGTATAG |
| **attL1-h NGLY1-int5-5HA-R** | CCTGCCTCAGTACTGAGAAGAGCTAGAGGTTAAAACAAGCACATG |
| **attL1-h NGLY1-int5-3HA-F** | CTAGCTCTTCTCAGTACTGAGGCAGGAGGATTGCTTGAACCTGGG |
| **attL1-h NGLY1-int5-3HA-R-b** | ACAAGAAAGCTGGGTAGATATCGGGCATTAGCACTAAAGTGGTATAGC |
| **hNgly1-SA-Exon6-F** | GTTTTAACCTCTAGCTCTTCTCAGTCTGACCTCTTCTCTTCCTCCCACAGATATAATAACCCTGAGAAACTTTTGG |
| **hNgly1-SA-Exon12-R** | GGGATGCCACCCGTTCAAAGGTCACTGAATTTTATAATTATC |
| **hGHpA-F** | CAGTGACCTTTGAACGGGTGGCATCCCTGTGACCCC |
| **hGHpA-R** | GGTTCAAGCAATCCTCCTGCCTCAGTATTTAAATAAGGACAGGGAAGGGAGCAGTG |
| **H592-mRuby-puro 5'-F** | GTGAACCACTGCTCCCTTCCCTGTCCTTATTTAAATGCATCAGATCTTTCCACTCAAAAC |
| **H592-mRuby-puro 5'-R** | GTTTTGAGTGGAAAGATCTGATGCATTTAAATAAGGACAGGGAAGGGAGCAGTGGTTCAC |
| **H592-mRuby-puro 3'-F** | GTAGAACATGAAATAACCTAGATCGATTTAAATACTGAGGCAGGAGGATTGCTTGAACC |
| **H592-mRuby-puro 3'-R** | GGTTCAAGCAATCCTCCTGCCTCAGTATTTAAATCGATCTAGGTTATTTCATGTTCTAC |
| **NGLY1-int5-SurF** | TTCCCCTGCCCAACCTAACCTACTC |
| **NGLY1-int5-SurR** | AGCTGAGCAGGAAAAGTGCTGCAGC |
| **NGLY1-testF1** | CAGGCTAAATCCAGCCGACATC |
| **NGLY1-qR3** | GGAAAGCTTCTTGCCCCATC |

**Supplementary Table S2**: List of antibodies used

| Target | Host species | Vendor | Catalog number |
|---|---|---|---|
| **β-actin** | Mouse IgG | Sigma-Aldrich | A5316 |
| **CD235** | Rabbit IgG | Thermo Fisher Scientific | PA5-27154 |
| **CD43** | Mouse IgG | Thermo Fisher Scientific | 14-0439-82 |
| **CD68** | Mouse IgG | Thermo Fisher Scientific | MA5-13324 |
| **CX3CR1** | Rabbit IgG | Thermo Fisher Scientific | PA128839 |
| **Cleaved caspase-3** | Rabbit IgG | Cell Signaling Technology | 9664T |
| **GAPDH** | Mouse IgG | Sigma-Aldrich | MAB374 |
| **IBA1** | Rabbit IgG | Wako | 019-19741 |
| **Ki67(8D5)** | Mouse IgG | Cell Signaling Technology | 9449S |
| **NGLY1** | Rabbit IgG | Millipore Sigma | HPA036825 |
| **NANOG** | Goat IgG | R&D Systems | AF1997 |
| **O-GlcNAc(RL2)** | Mouse IgG | Thermo Fisher Scientific | MA1-072 |
| **OGT** | Rabbit IgG | Cell Signaling Technology | 24083S |
| **OGA** | Rabbit IgG | Sigma-Aldrich | SAB4200311 |
| **OCT4** | Rabbit IgG | Cell Signaling Technology | 2840T |
| **p57 Kip2** | Rabbit IgG | Cell Signaling Technology | 2557T |
| **RUNX1** | Rabbit IgG | Abcam | ab23980 |
| **TMEM119** | Rabbit IgG | Thermo Fisher Scientific | PA-62505 |
| **TRA1-81** | Mouse IgM | BioLegend | 330702 |
| **FITC-affiniPure Donkey anti-Rabbit** | Donkey | Jackson Immuno Research | 711-095-152 |
| **Rhd-affiniPure Donkey anti-Rabbit** | Donkey | Jackson Immuno Research | 711-025-152 |
| **FITC-affiniPure Donkey anti-Mouse** | Donkey | Jackson Immuno Research | 715-095-150 |
| **Rhd-affiniPure Donkey anti-Mouse** | Donkey | Jackson Immuno Research | 715-025-150 |
| **FITC-affiniPure Donkey anti-Goat** | Donkey | Jackson Immuno Research | 705-095-003 |

**Supplementary Table S3:** qPCR primer sequences

| Gene Name | Forward (5'-3') | Reverse (5'-3') |
|---|---|---|
| *CCND1* | CGTCCATGCGGAAGATCG | AGCGGTCCAGGTAGTTCA |
| *CCNE1* | TCGCTCCAGGAAGAGGAA | CGCCGTCCTGTCGATTT |
| *CDK2* | GGTGGTGGCGCTTAAGAAA | GACATCCAGCAGCTTGACAATA |
| *CDKN1A* | CAGGTGGACCTGGAGACT | CGGATTAGGGCTTCCTCTTG |
| *CDKN1B* | GAGCAATGCGCAGGAATAAG | TCCACAGAACCGGCATTT |
| *CDKN1C* | GCGCGGCGATCAAGAAG | TTCTCAGGCGCTGATCTCTT |
| *COL3A1* | TGGTCTGCAAGGAATGCCTGGA | TCTTTCCCTGGGACACCATCAG |
| *CTGF* | GGCCAACCGCAAGATCG | TCTCCGCTGCGGTACAC |
| *FN1* | ACAACACCGAGGTGACTGAGAC | GGACACAACGATGCTTCCTGAG |
| *GAPDH* | ATGACATCAAGAAGGTGGTG | CATACCAGGAAATGAGCTTG |
| *HLA-DRA* | GCTCAGGAATCATGGGCTATC | CAAACATAAACTCGCCTGATTGG |
| *HLA-DRB1* | CCCACTGGCTTTGTCTGG | GCTCCGTCCCATTGAAGAAA |
| *IL-1B* | CTGATGGCCTAAACAGATGAA | TCGGAGATTCGTAGCTGGAT |
| *IL-6* | GACAAACAAATTCGGTACATCCTC | TTTCTGCCAGTGCCTCTTT |
| *MKI67* | GAGACGCCTGGTTACTATCAAA | ACACAACAGGAAGCTGGATAC |
| *PCNA* | AGATAACGCGGATACCTTGG | TCCATCAACTTCATTTCATAGTCTG |
| *RUNX1* | TCGGCTGAGCTGAGAAATG | TGTGAAGACAGTGATGGTCAG |
| *SPI1* | CGCCAAACGCACGAGTATTA | AGCTCTCGAACTCGCTGT |
| *TNF* | CCAGGGACCTCTCTCTAATCA | TCAGCTTGAGGGTTTGCTAC |
| *TREM2* | GAAGGTCCTGGTGGAGGT | ATCCTCGAAGCTCTCAGACTC |
| *TYROBP* | CTGCTCCTGCCTCTCCT | ACCGTAGAGCAACTGCAATC |

Supplementary Figure S1

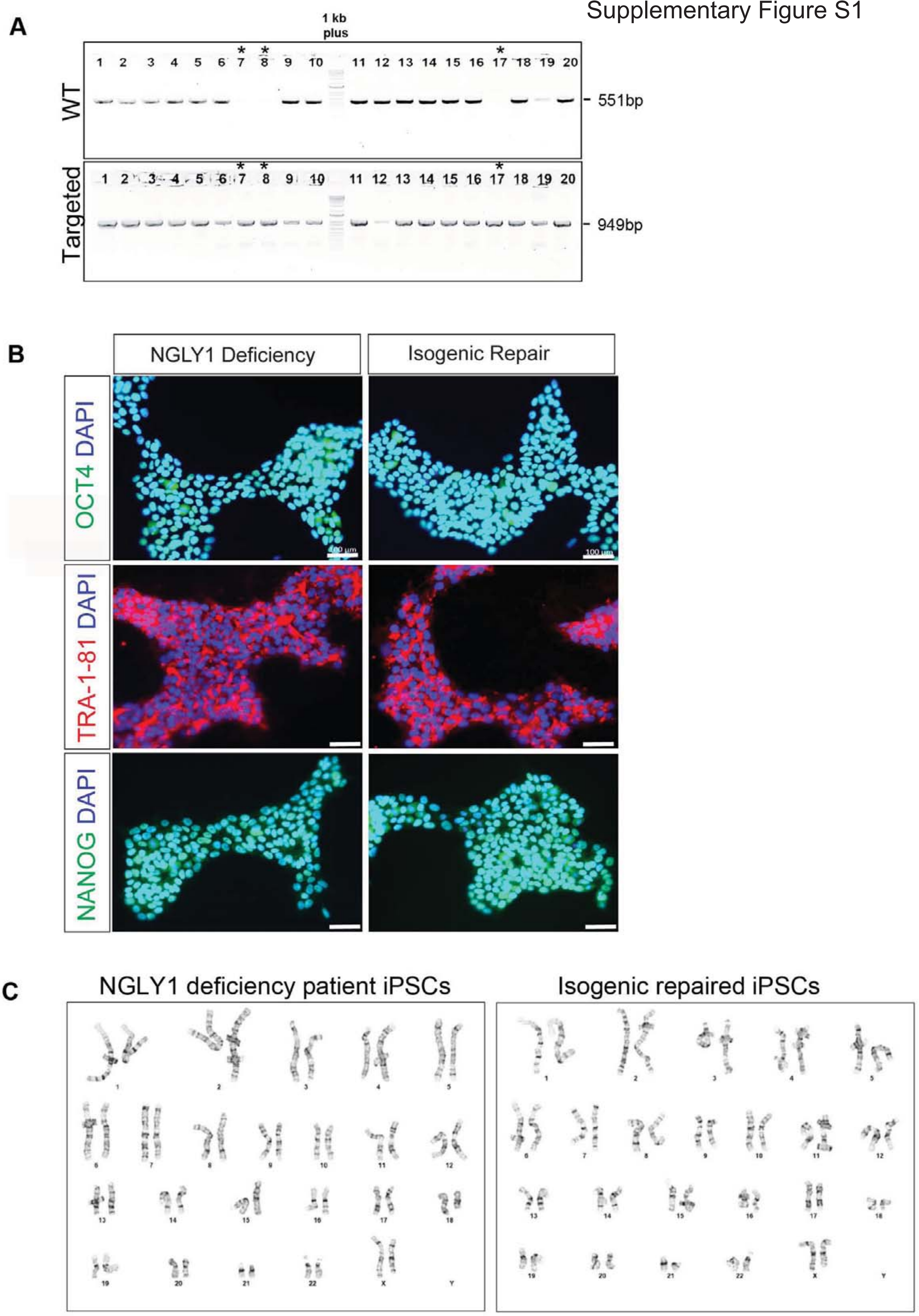

Supplementary Figure S2

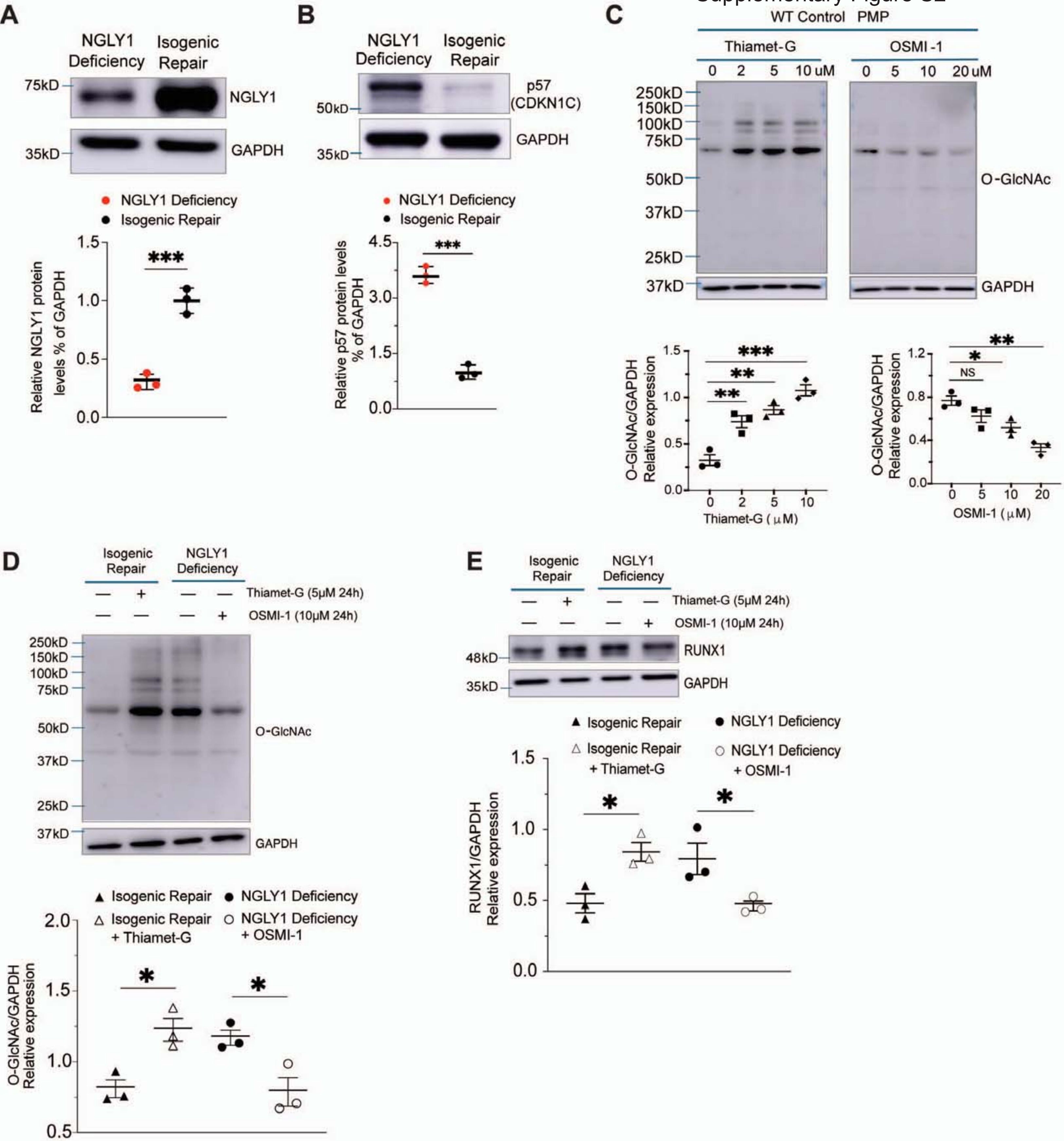

Supplementary Figure S3

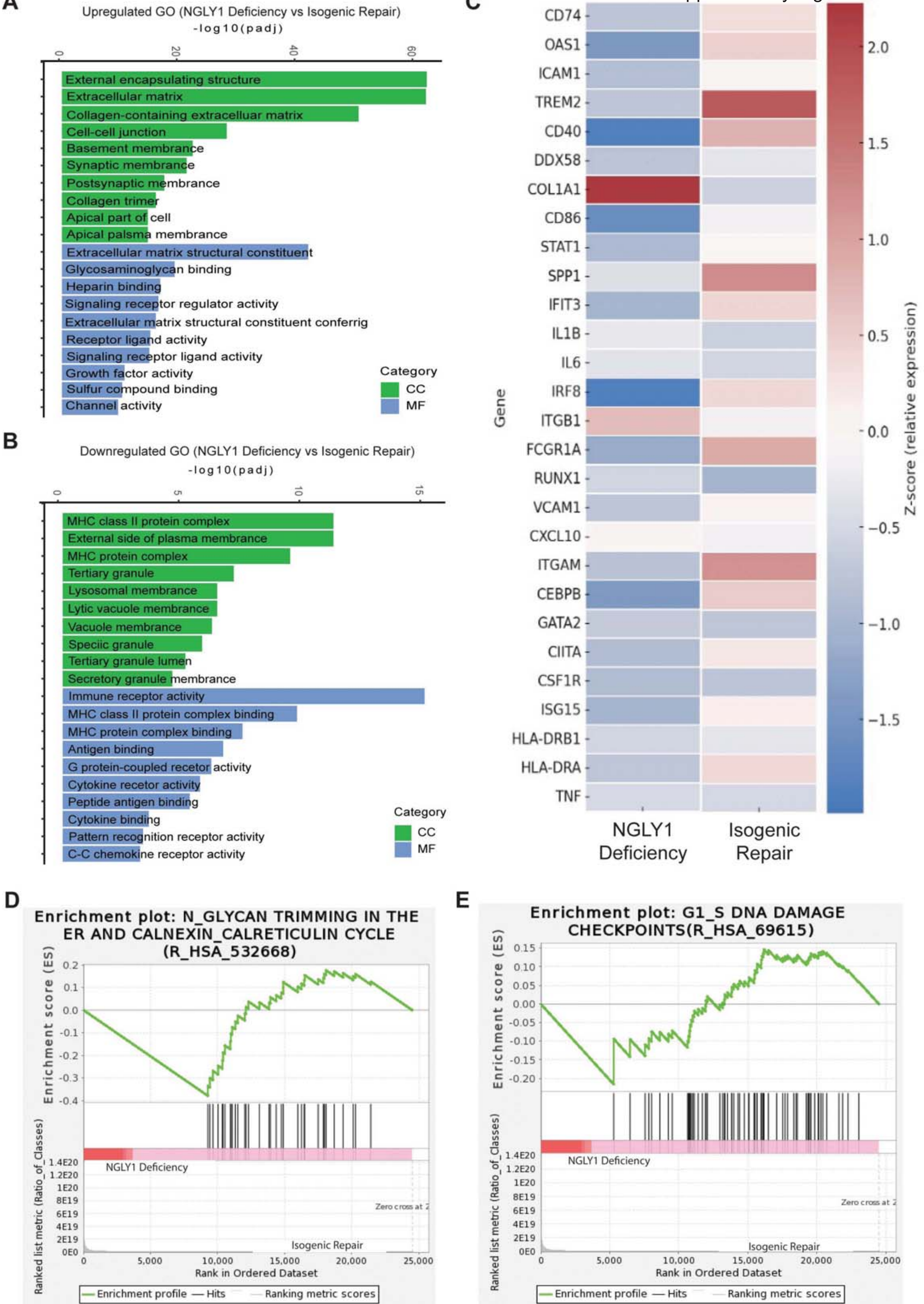

Supplementary Figure S4

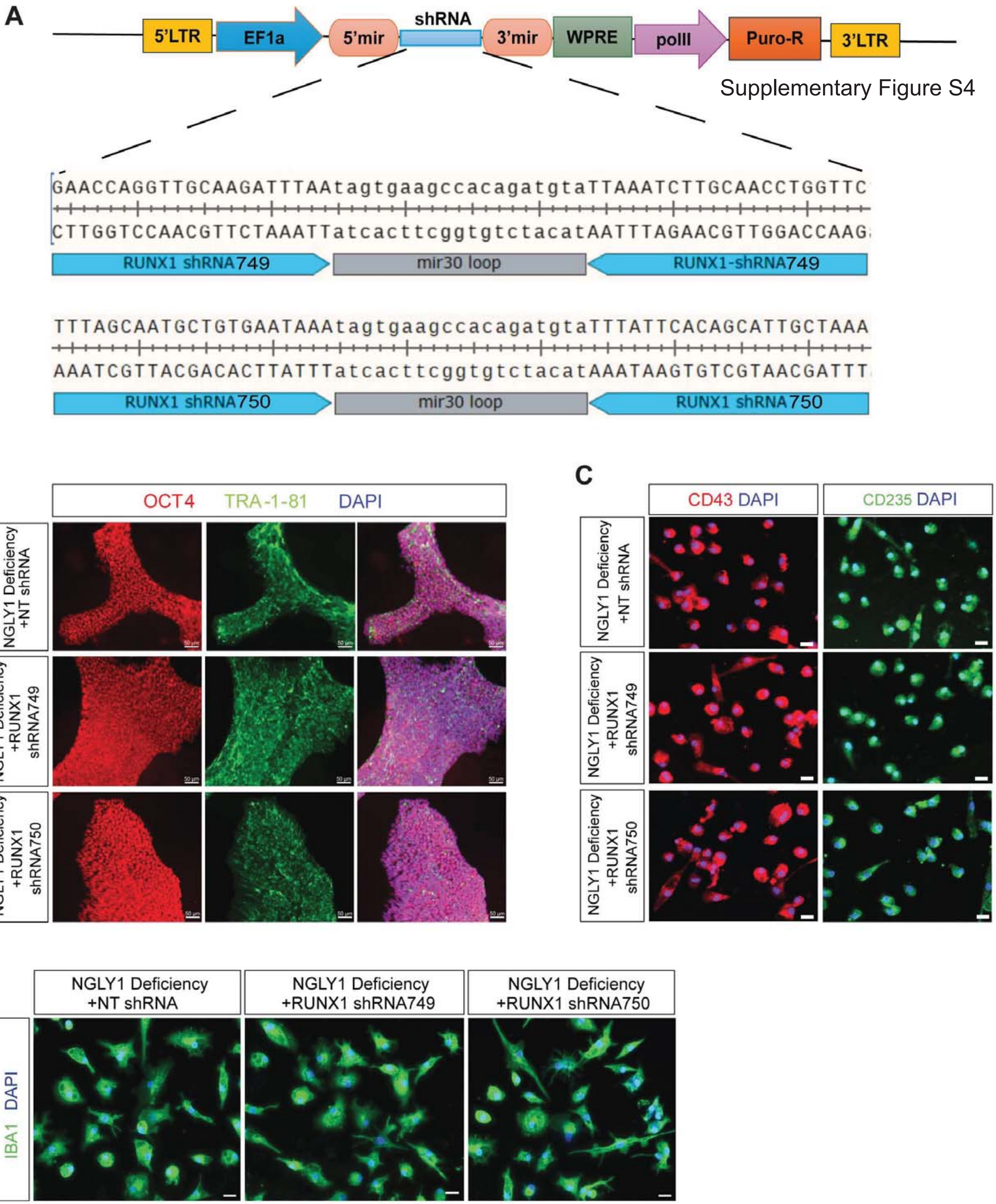
